# Explainable Artificial Intelligence for Cross-Dataset Generalizable Biomarker Discovery in Cardiovascular diseases (CVDs)

**DOI:** 10.64898/2026.07.23.740273

**Authors:** Ahtisham Fazeel Abbasi, Muhammad Sajjad, Sebastian Vollmer, Andreas Dengel, Muhammad Nabeel Asim

## Abstract

CVDs are heterogeneous, multifactorial disorders that remain the leading cause of global mortality from infancy to old age. It requires an early identification and treatment of risk factors to accelerate disease prevention and morbidity improvement. Advancements in transcriptomics technologies gives large pool of heterogenous gene expression data. The technical heterogeneity of gene expression data reduces ability to compare multiple cross-platform datasets at once. To bridge gap, we systematically evaluate three data harmonization techniques: Shambhala-2, TDM, and UPC to align heterogeneous data into a shared expression space while preserving biological signals. Our pipeline integrates 25 independent datasets comprising 983 samples across 23 distinct CVDs phenotypes from both RNA-seq and microarray platforms. The framework benchmarks 35 Machine learning (ML) and Deep learning (DL) classifiers, including Transformers and ResNets, across three data modalities such as RNA-seq, microarray hybridization and RNA-seq + microarray and multiple tissue types. To ensure clinical trustworthiness, we apply multiple Explainable artificial intelligence (XAI) methods, such as SHapley additive exPlanations (SHAP) and Integrated gradientss (IGs), and assess their reliability using quantitative metrics like Area over the perturbation curve (AOPC), Sensitivity, and Infidelity. Results indicate that Shambhala-2 provides superior harmonization by maximizing the biological signal-to-platform ratio. Evaluation of XAI methods reveals that Shapley-based approaches offer the highest stability for identifying influential genomic features in high-dimensional data. Functional enrichment and pathway analyses further confirmed the involvement of identified biomarkers in key cardiovascular processes, including inflammation, immune regulation, oxidative stress, and vascular remodeling. Collectively, this study provides a scalable and interpretable road-map that integrates XAI with cross-dataset biomarker discovery, supporting the transition toward precision cardiology.

## 1 Introduction

CVDs are a heterogeneous group of disorders of the heart and blood vessels [1] such as coronary artery disease, heart failure, cardiomyopathies, peripheral arterial disease, arrhythmias, and hypertension [2]. CVDs arise from a complex interplay of genetic, environmental, and molecular factors, encompassing non-modifiable risks such as age, sex, and genetic predisposition, as well as modifiable risks including hypertension, hyperglycemia, dyslipidemia, smoking, unhealthy diet, obesity, and physical inactivity [3–5]. Globally, CVDs remain the leading cause of death, accounting for nearly one-third of all deaths each year [6], with ischemic heart disease and stroke contributing to the majority of cases [7]. In particular, 80% of these deaths occur in low- and midDLe-income countries [7, 8], where limited access to preventive care, early diagnosis, and effective treatment results in late disease detection and increased premature mortality [9]. Beyond mortality, CVDs place a heavy burden on healthcare systems through high hospitalization, medication, and long-term care costs [10].

Conventionally, CVDs are diagnosed using a combination of clinical assessment, laboratory testing, and imaging studies [11]. Initial evaluation involves medical and family history, physical examination, and cardiac auscultation [12, 13], followed by laboratory measurements of cardiac biomarkers (e.g., troponins, CK-MB, myoglobin) and risk factors such as cholesterol and blood glucose [14], alongside imaging including echocardiography (ECG), computed tomography (CT), and magnetic resonance imaging (MRI) [15, 16]. Although these approaches are effective for detecting established disease, they often lack sensitivity for early or subclinical pathological changes [17], rely on biomarkers with limited disease specificity [18], and involve costly or invasive procedures [18]. Consequently, these clinical, laboratory, and imaging-based [19] approaches have limited ability to capture the underlying molecular heterogeneity of CVDs and to reliably detect early disease states [20]. This motivates the need for robust and generalizable molecular-level biomarkers, supported by computational frameworks that integrate high-throughput data with computational tools, to enable earlier, more precise, and scalable CVDs detection and characterization [21, 22].

Building on the limitations of conventional diagnostic [23] and statistical methods [24] in identifying biomarkers in CVDs, the integration of Artificial intelligence (AI) with high-throughput omics data has emerged as a promising framework for improved diagnosis and biomarker discovery [25]. To the best of our knowledge, only 63 studies have investigated CVDs diagnosis and biomarker discovery. Among these, 45 studies utilize transcriptomic data for disease classification in conjunction with biomarker discovery. Notably, the majority of existing work focuses on single-disease classification, with acute myocardial infarction (14) [26–38], heart failure (7) [39–45], and atherosclerosis (6) [46–51] being the most extensively studied. These are followed by ischemic stroke (3) [52–54], hypertrophic cardiomyopathy (3) [55–57], and general CVDs classification (3) [58–60]. Other conditions namely, atrial fibrillation (2) [61, 62], coronary artery disease (2) [63, 64], congenital heart disease [65], acute aortic dissection [66], kawasaki disease [67], rheumatoid arthritis [68], and cardiovascular calcification [69], remain comparatively underexplored. Methodologically, the literature primarily focuses on classical ML approaches, including Support vector machines (SVMs), Random forests (RFs), K nearest neighbors (KNNs), Gradient boosting (GB), and decision trees (DTs), frequently combined with feature selection techniques such as boruta and recursive feature elimination (RFE) [38, 53, 69]. In contrast, only a small subset of studies has explored DL–based models [29, 30, 41, 48, 49, 52, 67]. These studies mainly employed neural network–based architectures, including artificial neural networks (ANNs) [48, 52, 57] and Convolutional neural networks (CNNs) [41, 49], often combined with feature selection or ensemble techniques to enhance classification performance.

Existing studies on CVDs diagnosis and biomarker discovery suffer from several interconnected limitations. For instance, most studies are restricted to single-disease i.e., binary classification settings, thereby failing to capture shared and disease-specific molecular patterns across multiple CVDs conditions [70, 71]. Moreover, existing studies rarely investigate data harmonization and integration strategies to combine heterogeneous transcriptomic datasets across different studies or platforms, which limits the discovery of generalizable biomarkers in multiclass classification settings [72, 73]. In addition, classical feature selection techniques are more commonly utilized, while explainable artificial intelligence (XAI) approaches largely overlooked, resulting in limited biological interpretability and mechanistic insight [74]. In addition, most studies do not explicitly evaluate the quality, robustness, or stability of selected biomarkers using dedicated quantitative evaluation metrics, relying instead on downstream classification performance as a proxy for biomarker validity [75]. Finally, existing studies offer minimal performance comparison against transformer-based models, despite their demonstrated potential for modeling high-dimensional omics data [76, 77]. Collectively, these shortcomings highlight a critical gap in the current literature: the absence of a harmonized, multi-CVDs molecular dataset and framework that enables robust multiclass diagnosis, generalizable biomarker discovery, and explainable, state-of-the-art model evaluation.

### 1.1 Summary

CVDs transcriptomic datasets (mRNA) i.e,. RNA-seq and microarray, are highly dispersed across public repositories and are typically analyzed in isolation, with most studies developing disease-specific predictors tailored to a single cohort or condition. This dispersion, combined with substantial heterogeneity in platforms, sample sizes, preprocessing protocols, and disease phenotypes, limits the development of robust and generalizable CVDs biomarkers. To address these challenges, we propose a unified large-scale analytical framework that integrates diverse CVDs transcriptomic (mRNA) datasets and systematically evaluates predictive modeling and biomarker discovery across multiple disease categories.

In the first step, transcriptomic datasets (mRNA) are collected from public resources, including the Gene Expression Omnibus (GEO) [78] and CVDs-focused repositories such as CVDs Atlas [79]. The curated datasets generated from multiple CVDs, tissues, and experimental protocols. Standardized preprocessing steps are applied to all datasets, including quality control, gene annotation, gene filtering, and normalization, to ensure consistency across studies while preserving biologically meaningful variation.

In the subsequent step, technical variability arising from differences in experimental protocols and tissue sources is addressed through data harmonization using Shambhala-2 [80], Universal exPression Code (UPC) [81] and Training Distribution Matching (TDM) [82]. These harmonization strategies aim to project heterogeneous datasets into a shared expression space while maintaining relative gene expression patterns critical for downstream analyses and biomarker discovery. The effectiveness of data harmonization is evaluated using the Watermelon multi-selection algorithm [83] by examining the stability of gene expression signals across harmonized datasets.

Following harmonization, both ML and DL models are trained for CVDs diagnosis. The evaluated models capture diverse inductive biases, ranging from classical ML approaches to neural network–based architectures, enabling a comprehensive assessment of predictive performance across CVDs. Model evaluation is conducted using standardized metrics to ensure fair comparison and to quantify robustness across heterogeneous cohorts.

To enhance interpretability and facilitate biomarker discovery, XAI methods are applied to trained ML and DL models, including SHAP [84], Local interpretable model-agnostic explanations (LIME) [85], DL important features (DeepLIFT) [86], IGs [87], and Layer-wise relevance propagation (LRP) (LRP) [88]. These methods provide complementary perspectives on feature importance and attribution, enabling the identification of genes that drive model predictions across different CVDs phenotypes. The quality and reliability of the explanations is further assessed using quantitative explanation evaluation metrics, including area over the perturbation curve (AOPC) [89], maximum sensitivity, and infidelity [90], to ensure that the biomarkers identified are not only predictive but also robust and generalizable.

Figure 1 illustrates an overview of the proposed CVDs analysis pipeline, highlighting the complete workflow from dataset collection and preprocessing to harmonization, predictive modeling, explainability, evaluation, and downstream analyses.

**Figure 1:**
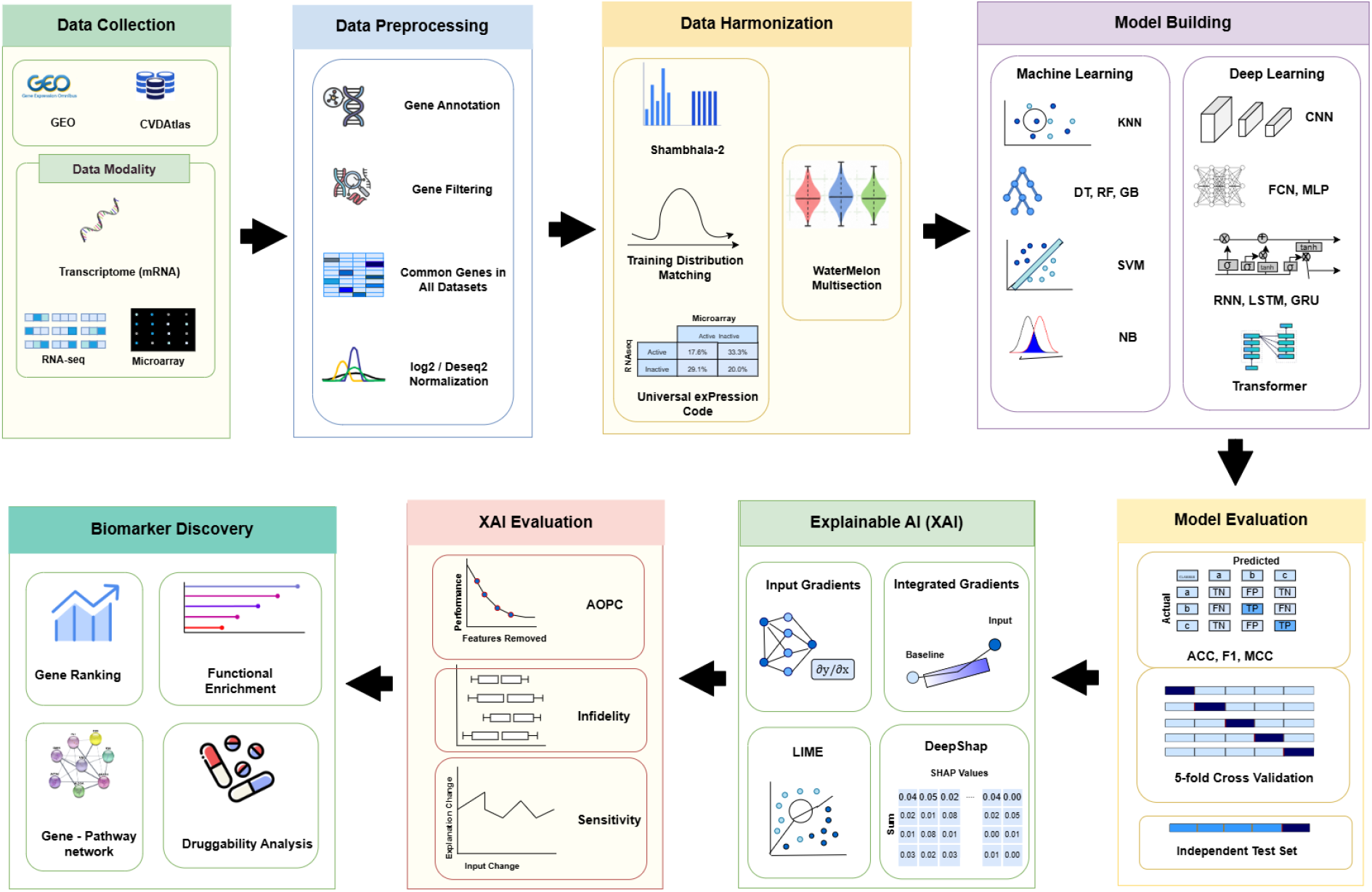
An end-to-end pipeline for heterogeneous data harmonization and CVDs prediction, integrating ML/DL models with XAI techniques.

Detailed descriptions of dataset collection and preprocessing are provided in section 1.2. Data harmonization strategies are described in section 1.3, while classification models are presented in sections 1.4. Explainability methods and explanation quality metrics are discussed in sections 1.5 and 1.6. Together, this framework enables a systematic, scalable, and reproducible analysis of CVDs transcriptomic data, supporting robust disease classification and interpretable biomarker discovery across heterogeneous cohorts.

### 1.2 CVDs Benchmark Dataset

Transcriptomic studies of CVDs (CVDs) often rely on publicly available gene expression datasets. However, these data are distributed across multiple repositories, experimental platforms, and disease-specific cohorts, which complicates their integration and systematic reuse. Consequently, many existing studies focus on single-disease datasets, limiting the ability of computational models to identify both shared molecular signatures and disease-specific markers across different CVDs conditions. To address this limitation, we assembled a multi-disease, multi-platform transcriptomic dataset by integrating publicly available gene expression data from the Gene Expression Omnibus (GEO) and CVDs Atlas.

In this study, dataset selection for CVDs research is guided by a set of predefined criteria to ensure data consistency and disease relevance across two distinct transcriptomic modalities, namely microarray and RNA-seq. The inclusion criteria for each dataset are defined as follows: (i) the dataset must originate from human subjects, (ii) it must be directly relevant to a CVDs, (iii) it must contain coding gene (mRNA) expression data. (iv) it must contain confirmed diseased status with available meta-data, and (v) the CVDs should not be associated with other diseases or viral infections. Datasets with incomplete or unclear disease annotations are excluded. Datasets grouped according to tissue type, and only datasets from the same tissue integrated together. This approach helped preserve tissue-specific gene expression patterns and prevent confounding biological variation arising from differences in tissue composition. Control samples are removed before harmonization so that the final data matrices contain only diseased samples, ensuring that downstream normalization and classification models focus on clinically relevant molecular variation between CVDs rather than comparisons with healthy samples.

Table 1 provides an overview of 12 RNAseq datasets comprising 515 samples across eleven CVDs, generated using eight distinct sequencing platforms and obtained from GEO and CVDs Atlas. These datasets span three tissue types: blood, left ventricle, and heart. Blood tissue datasets represent five CVDs that can be assessed through peripheral or circulating gene expression signals, including coronary artery disease (n = 27, GPL23934), acute myocardial infarction, comprising both ST-elevation myocardial infarction (STEMI) and non-ST-elevation myocardial infarction (NSTEMI) cases (n = 30, GPL16288), rheumatic heart disease (n = 14, GPL20795), dilated cardiomyopathy (n = 7, GPL20301), and acute coronary syndrome, including patients with and without aortic valve sclerosis (AVSc) (n = 101, GPL16288). Left ventricle datasets represent diseases involving direct myocardial pathology, including dilated cardiomyopathy (n = 161, n = 37, GPL16791), peripartum cardiomyopathy (n=6, GPL16791), hypertrophic cardiomyopathy (n=28, GPL16791), idiopathic dilated cardiomyopathy (n=15, n=7 across GPL16791 and GPL20301), ischemic cardiomyopathy (n=13, GPL1679), and myocardial ischemia (n = 8, GPL18573). Heart tissue datasets include ischemic cardiomyopathy (n = 14, GPL9115) and heart failure (n = 47, GPL11154). Each RNAseq dataset contains 39,376 features corresponding to genes shared across datasets within the same tissue group. RNAseq preprocessing is performed in R (v4.5.2) using the GEOquery [91], dplyr [92], AnnotationDbi [93], and org.Hs.eg.db [94] packages. Within each tissue group, the feature space is restricted to the intersection of genes common across all contributing datasets. Genes lacking valid HGNC symbols are removed, and duplicate gene entries are resolved by retaining a single representative per gene.

**Table 1:**
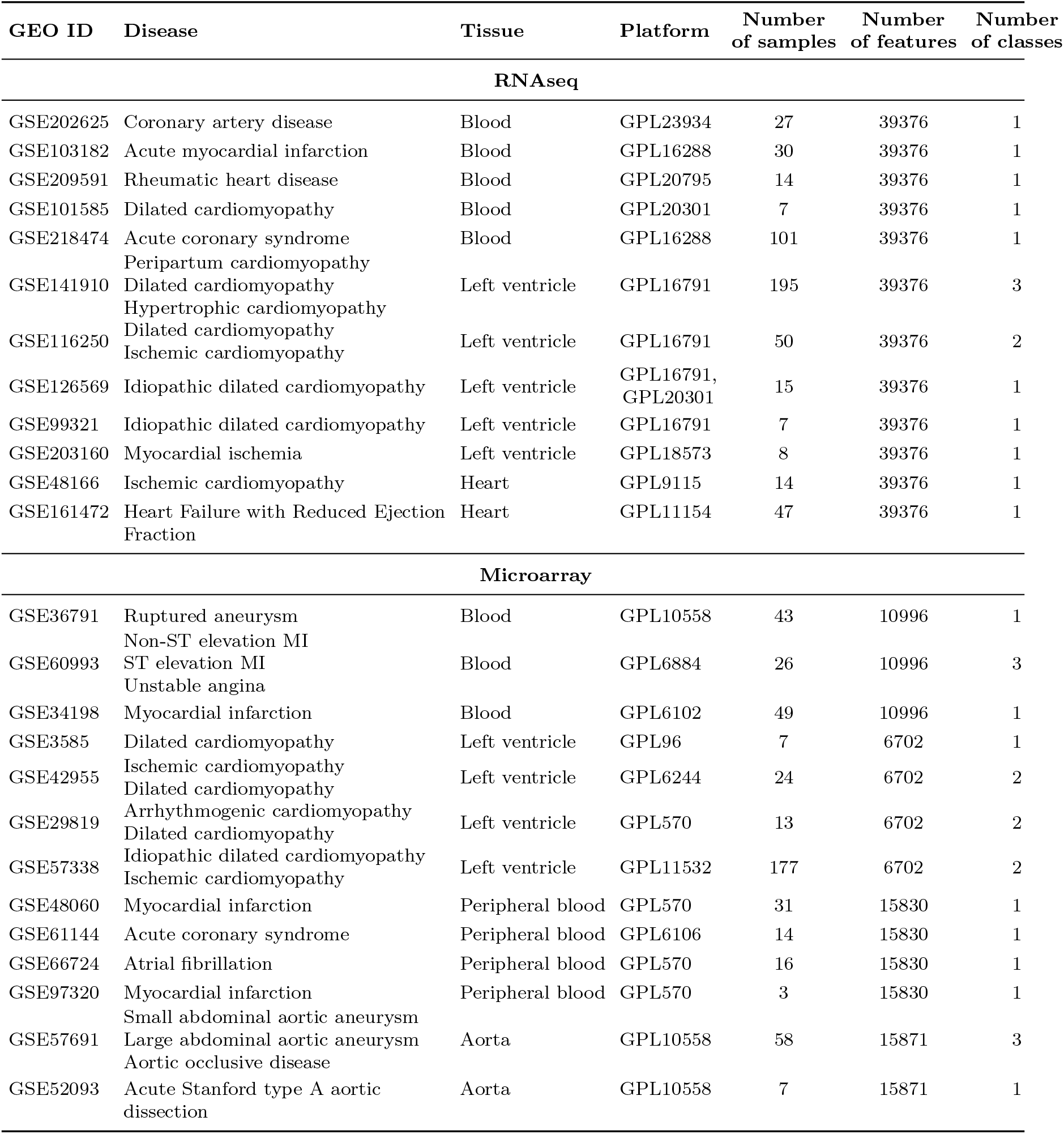
Gene Expression Data Statistics for Harmonization.

Microarray datasets are obtained exclusively from Affymetrix array technologies across eight platforms and organized into four tissue groups: left ventricle, blood, peripheral blood, and aorta. Left ventricle datasets include four CVDs, namely dilated cardiomyopathy (n = 7, GPL96; n = 12, GPL6244; n = 6, GPL570), ischemic cardiomyopathy (n = 12, GPL6244; n = 95, GPL11532), arrhythmogenic cardiomyopathy (n = 7, GPL570), and idiopathic dilated cardiomyopathy (n = 82, GPL11532). Blood datasets include ruptured aneurysm (n = 43, GPL10558), non-ST-elevation myocardial infarction (n = 10, GPL6884), ST-elevation myocardial infarction (n = 7, GPL6884), unstable angina (n = 9, GPL6884), and myocardial infarction (n = 49, GPL6102). Peripheral blood datasets include myocardial infarction (n = 34, GPL570), atrial fibrillation (n = 16, GPL570), and acute coronary syndrome (n = 14, GPL6106). A set of samples was taken from peripheral blood only, to reduce biological and technical variability and analyzed separately. The blood/whole blood/peripheral blood data combined is expected to maximize sample size and include more blood related transcriptomic patterns, but these sample types may differ in cellular composition and pre-processing characteristics, which can introduce heterogeneity. Hence, the peripheral blood-only set is used as a more uniform set for validation of consistency. Aorta datasets include small and large abdominal aortic aneurysm, aortic occlusive disease (n = 58) and acute Stanford type A aortic dissection (n = 7), both generated on GPL10558. Log2 transformation is applied when expression values indicate non-transformed data, and genes with missing values are removed to ensure complete matrices. Finally, features are restricted to the intersection of genes shared across datasets within each tissue group to maintain cross-platform consistency.

Overall, the benchmark includes 983 samples covering 23 cardiovascular disease phenotypes from 25 independent datasets. This multi-disease composition allows the framework to perform multiclass classification across a clinically relevant range of CVDs, including cardiomyopathies, ischemic syndromes, arrhythmias, and vascular inflammatory diseases, while maintaining meaningful molecular differences between diseases. Inclusion of both RNAseq and microarray datasets further enables cross-platform validation and tests the robustness of harmonization strategies, as described in Section 1.3.

### 1.3 Data Harmonization

Data originating from heterogeneous sources, platforms, and tissue types cannot be directly combined or used in AI models due to systematic technical variations, batch effects, and distributional shifts that can introduce spurious patterns and degrade model generalizability [95, 96]. Such heterogeneity may lead to biased predictions and reduced robustness [97]. Therefore, data harmonization is a critical step to align datasets into a common representation while preserving underlying biological signals [98]. In this study, we utilize three widely used harmonization approaches, TDM [82], Shambhala-2 [80], and UPC [81], to mitigate cross-dataset variability. These approaches are described in detail in the following subsections.

- ***Training Distribution Matching (TDM):*** AI models typically assume that the training and test datasets are drawn from the same underlying distribution. When this assumption is violated, model performance declines, leading to a problem known as dataset shift [99]. To address this, TDM [82] is developed as a data harmonization and normalization approach. TDM aligns the feature-wise distribution of a target dataset to match that of a reference training dataset, thereby reducing distributional differences and platform-specific variability. In gene expression analysis, this helps models trained on RNAseq data from one source to generalize reliably to data from other sources. TDM also enables cross-platform transfer by transforming the target dataset to match the training distribution, allowing models trained on microarray data to be applied to RNAseq datasets, or vice versa. By bringing heterogeneous datasets into a common statistical space while preserving biologically meaningful signals, TDM facilitates the integration of multi-source data for downstream ML and AI analyses. The working paradigm of TDM is comprised of four main steps designed to harmonize RNAseq expression profiles across different tissues and sequencing platforms. First, RNAseq expression values in the reference (training) dataset are log_2_-transformed to stabilize variance and ensure comparability across datasets, ensuring consistency with microarray data, which are typically available in log_2_-transformed form. In the next step, the distribution of the training dataset is summarized using the first quartile (Q1), third quartile (Q3), interquartile range (IQR), and the minimum and maximum expression values. These statistics capture the central tendency and spread of the expression values while remaining resilient to platform-specific noise and outliers. Based on these training statistics, two scaling ratios are computed to describe the relationship between the central data distribution and the upper and lower distribution tails of the training data:

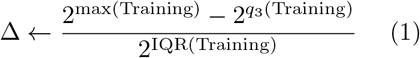

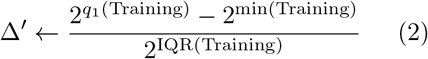 These ratios are then transferred to the target (testing) dataset to estimate dataset-specific upper and lower expression bounds using its own quartiles and IQR.

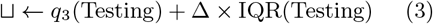

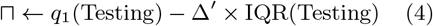 Expression values in the target dataset are subsequently clipped to these bounds to control extreme values arising from technical variability across tissues and sequencing machines, while ensuring all values remain non-negative. Finally, the clipped expression values are rescaled to match the dynamic range of the training dataset, which allows both datasets to have comparable distributional properties.

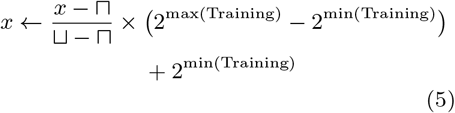 By applying TDM, heterogeneous RNAseq datasets are aligned into a common distributional space, reducing platform-induced biases while preserving biologically meaningful variation, thereby enabling reliable downstream ML and AI modeling across multi-source CVDs datasets.
- ***Shambhala-2:*** Shambhala-2 [80] is an extension of the previous Shambhala-1 [100] framework, developed to address batch effects and platform-specific biases in gene expression data. Shambhala-1 was designed to process multiple samples originating from different experimental platforms, allowing newly harmonized data to be added to a pre-calculated reference pool without requiring recomputation of the entire dataset. Shambhala-2 combines the advantages of Shambhala-1 with the CuBlock approach [101], which applies piecewise cubic transformations for harmonization, thereby improving harmonization across experimental platforms. In the Shambhala methods [80, 100], the core concept is the generation of a universal output format, ensuring that any dataset processed through Shambhala remains directly comparable to all others. This is achieved by transforming gene expression profiles into a pre-defined reference format, referred to as the reference definitive dataset *Q*. During harmonization, newly added expression data are adjusted to match the statistical characteristics of this fixed reference dataset *Q*, ensuring consistent distributional properties across datasets processed at different times and from different platforms. Shambhala-2 [80] processes gene expression data through a multi-step harmonization pipeline. The procedure begins with an input expression matrix, which can originate from any platform, tissue, or experimental protocol. The raw dataset is first combined with an auxiliary calibration dataset *P* to perform quantile normalization, resulting in *P* → *P*^*′*^. This step ensures that the datasets are statistically comparable by aligning their gene expression distributions. Specifically, genes within each sample are ranked, reference quantiles are computed from *P*, and the original values are replaced by the rank means, after which the original gene order is restored. In the next step, the quantile-normalized dataset *P*^*′*^ is processed using the CuBlock method (*P* ^*′*^ → *P* ^*′′*^), which performs platform-independent normalization by reshaping local gene expression distributions within each sample. First, genes are partitioned into multiple blocks using k-means clustering to capture local structure in the expression space. Within each block, expression values are standardized using z-score normalization and then subjected to rank-based mapping to predefined target values, followed by density shaping. A cubic polynomial is subsequently fitted to the transformed values and applied to the original z-scores to adjust the distribution. This process is repeated across all blocks and samples, with the clustering performed multiple times and the results averaged to ensure stability. Finally, the normalized dataset *P* ^*′′*^ is genewise rescaled using the mean 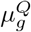 and standard deviation 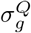 from the reference definitive dataset *Q*. This ensures that any dataset processed through Shambhala-2 retains consistent distributional properties and remains directly comparable to previously harmonized datasets.
- ***UPC:*** UPC [81] is a probabilistic barcoding method that estimates the transcriptional activity of genes within a sample. Its goal is to classify each gene as either active or inactive, allowing raw gene expression values from different platforms to be interpreted in a comparable framework. The UPC approach begins with a raw expression matrix, which can originate from microarrays (single- or two-color) or RNAseq experiments. Each gene’s observed expression is assumed to arise from one of two distributions: a background distribution representing inactive genes, and a background-plus-signal distribution representing active genes. An unobserved indicator variable Δ_*i*_ is introduced for each gene *i*, taking the value 1 if the gene is active and 0 if inactive. For each gene, the background distribution is estimated using statistical models that account for relevant features such as probe composition, nucleotide content, gene length, or other platform-specific factors. Genes with similar features are expected to have similar background distributions, while active genes are assumed to exhibit an additional signal on top of this background. UPC [81] employs a mixture model combining the background and background-plus-signal components. The proportion of active genes (*π*) and the parameters of each distribution (*θ*_1_ for background, *θ*_2_ for background-plus-signal) are estimated using the expectation–maximization (EM) algorithm. The EM algorithm iteratively assigns probabilistic memberships to each gene (expectation step) and updates the model parameters using these probabilities (maximization step) until convergence. After convergence of the EM algorithm, the UPC value for each gene *i*, denoted *P*_*i*_, is calculated as the posterior probability of the gene being active:

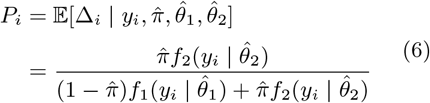

where *f*_1_ and *f*_2_ are the density functions of the background and background-plus-signal distributions, respectively. This transforms raw expression values into probabilities between 0 and 1, representing gene activity. The final output is a matrix of UPC values for all genes in the sample. These values are directly comparable across samples and platforms, providing a platform-independent measure of transcriptional activity suitable for downstream analyses.

### 1.4 Classifiers

The performance of AI classifiers for CVDs classification across the proposed benchmark datasets is assessed by exploring the potential of 35 distinct ML (15) and DL (20) classifiers. These classifiers include tree-based, distance-based, and neural network models, which provide diverse algorithmic coverage and maintain manageable computation across all datasets. The selected classifiers are consistent with those most frequently used in recent CVDs classification studies, allowing direct comparability of results [102, 103].

Naive Bayes (NB) is a probabilistic generative classifier that applies Bayes’ theorem to estimate the likelihood of a sample belonging to a given class based on observed feature patterns. By assuming conditional independence among features, NB computes class probabilities efficiently and assigns labels according to the most probable class [104]. Logistic Regression (LR) [105] is a discriminative classifier that focuses directly on learning the decision boundary between classes. LR learns a set of feature weights and an intercept term, where each weight quantifies the contribution of a feature to class separation. The weighted combination of features is transformed using a logistic (sigmoid) function to yield probabilistic predictions.

Tree-based classifiers, including Decision Trees (DTs) [106] and ensemble methods such as RFs [107] and boosting-based methods, are non-parametric classifiers capable of learning complex, non-linear decision boundaries. DTs recursively partition the feature space using nested if–then–else rules, where splits are determined by criteria such as Gini impurity or information gain. RFs extend this framework by constructing multiple decision trees on bootstrapped samples of the training data and aggregating their predictions via majority voting. Boosting methods, including AdaBoost (AB) [108], GB [109], Histogram-Based Gradient Boosting (HGB), and optimized frameworks such as Extreme Gradient boosting (XGB) [110], Light Gradient Boosting Machine (LGBM) [111], and Categorical Boosting (CB) [112], iteratively train weak learners while emphasizing previously misclassified samples. These approaches minimize loss functions through gradient-based optimization, incorporate regularization and efficient tree growth strategies, and are particularly effective for heterogeneous and structured biomedical data.

Distance-based classifiers such as KNN [113] assign class labels based on the majority vote of the k closest training instances in feature space, typically measured using Euclidean distance. The resulting decision boundary can take a flexible, non-linear shape that adapts to the distribution of the training data. Quadratic Discriminant Analysis (QDA) [114, 115] is a probabilistic classifier that assumes class-specific Gaussian distributions with distinct covariance matrices, resulting in quadratic decision boundaries.

SVMs [116, 117] are large-margin classifiers that learn optimal decision boundaries by maximizing the margin between classes. For non-linearly separable data, SVMs use kernel functions to project samples into higher-dimensional feature spaces, enabling linear separation without explicitly computing the transformation. This kernel trick allows SVMs to model complex decision boundaries while maintaining computational efficiency.

DL classifiers capture complex patterns in high-dimensional gene expression data that are difficult to model using traditional ML classifiers. Multi-layer perceptrons (MLPs)[118] are fully connected, feed-forward networks in which input features pass through multiple layers of neurons, allowing the model to learn non-linear relationships through iterative weight updates during training.

CNNs [119] learn useful patterns by applying small filters to the input data, followed by pooling operations that reduce data size while keeping important information [120]. ResNets [121] improve CNN performance by adding skip connections that allow information to pass directly across layers, making it easier to train deeper models [122]. Densely connected convolutional networks (DenseNets) [123] create direct connections between all layers within a dense block. This connectivity lets each layer receive features from all preceding layers, enabling feature reuse. This design helps DenseNet variants, including DenseNet121, DenseNet161, and DenseNet169, to achieve strong performance with fewer parameters [124].

RNN-based models [122, 125] are designed to hanDLe ordered or sequential data by retaining information from previous inputs. However, standard RNNs have difficulty learning long-term patterns [126]. Long short-term memory (LSTMs) [127] address this issue by using gating mechanisms that control what information is stored or discarded over time. Gated recurrent units (GRUs) [128, 129] offer a simpler alternative to LSTMs with fewer components, while still effectively learning sequential dependencies. Hybrid models such as CNN-LSTM and CNN-GRU combine convolutional layers with recurrent layers, enabling them to learn both local feature patterns and sequential relationships [130].

The DeepGene Transformer [131] is an attention-based model developed specifically for gene expression analysis. It combines self-attention with one-dimensional convolutional layers to identify both global and local relationships among genes. By focusing more on informative genes, the model can learn meaningful biological patterns without requiring manual feature selection, making it well suited for large-scale gene expression studies.

### 1.5 XAI

To move beyond black-box predictions and enable biological interpretation, we applied XAI techniques to quantify the contribution of individual genes to model decisions. This approach allows the identification of disease-specific and shared molecular signatures, facilitating the discovery of candidate biomarkers with potential clinical relevance. XAI encompasses a collection of methods designed to improve the transparency, interpretability, and trustworthiness of ML and DL models by clarifying how predictions are made [132].

This is particularly important in biomedical and clinical domains, where understanding the reasoning behind a model’s decision is as critical as predictive performance [133]. XAI methods aim to answer key questions such as which features are important, why a specific prediction is made, and whether the learned patterns are biologically meaningful or clinically credible.

- **Taxonomy of XAI Methods:** Multiple taxonomies have been proposed to to categorize AI interpretability models, but most classify these methods into four main categories: scope-based, complexity-based, model-based, and methodology-based. Scope-based methods focus on the level at which explanations are provided. Local methods explain individual predictions by showing why a model produced a specific output for a given input, whereas global methods describe the model’s overall behavior across the entire dataset. LIME is widely used for local explanations, and SHAP can provide both local and global insights. The complexity-based taxonomy differentiates methods based on when and how interpretability is achieved, separating intrinsic (ante-hoc) approaches, which rely on inherently interpretable models such as linear models and decision trees, from post-hoc approaches, which are applied after training to explain complex blackbox models, including techniques such as LIME, SHAP, and saliency-based methods. The model-based taxonomy categorizes methods according to their dependence on the underlying model, distinguishing model-agnostic approaches, which can be applied to any predictive model given access to inputs and outputs (e.g., LIME and SHAP), from model-specific approaches, which exploit internal model structures such as gradients, activations, or layer-wise computations, including methods like IGs, DeepLIFT, and LRP. Finally, the methodology-based taxonomy groups methods based on how explanations are generated, distinguishing perturbation-based approaches, which assess feature importance by systematically modifying inputs and observing output variations (e.g., LIME, SHAP, and RISE), from gradient-based approaches, which utilize backpropagation and gradient information to attribute relevance scores to input features, such as saliency maps, Grad-CAM, IGs, DeepLIFT, and LRP. In this work, we adopt a structured selection of XAI methods aligned with the major taxonomic categories to ensure a comprehensive and balanced evaluation of explainability techniques. From the scope-based perspective, both local and global explainability methods are considered, where LIME is employed for local interpretability to provide instance-level insights into individual predictions, while DeepSHAP is utilized for its capability to generate both local explanations and aggregated global feature attributions. From a complexity-based standpoint, we focus on post-hoc methods, as they are well-suited for explaining high-complexity DL models without altering their architecture; accordingly, IG and Input Gradients are selected due to their effectiveness in attributing predictions in neural networks. From the model-based perspective, we incorporate both model-agnostic and model-specific approaches to ensure methodological diversity, where LIME and DeepSHAP represent model-agnostic techniques applicable across different architectures, while IGs and Input Gradients serve as model-specific methods that leverage internal network information such as gradients. Finally, from the methodology-based viewpoint, both perturbation-based and gradient-based techniques are included, with LIME and DeepSHAP representing perturbation-based approaches, and IGs alongside Input Gradients representing gradient-based attribution methods, thereby enabling a comparative analysis of different explanation strategies and their impact on interpretability.
- **IG:** Deep neural networks (DNNs), despite their high performance, often behave like black boxes, making it difficult to interpret how individual inputs lead to specific outputs [134]. This lack of transparency limits the ability of human experts to understand the reasoning behind each prediction, restricting the practical application of DL models in domains that require trust and accountability. Feature attribution in neural networks helps address this issue by identifying which input features are important for a model’s final prediction, providing insight into model behavior, and ensuring transparency. However, most existing attribution methods fail to address two common axioms, sensitivity and implementation variation [135]. Sensitivity requires that if a change in an input causes a different prediction, the method should assign non-zero attribution to that input. Implementation variation reflects differences in attributions across functionally equivalent models. Among attribution methods, IG is particularly notable because it overcomes these issues, providing more consistent and reliable explanations for model predictions. IG [87] combines the principles of gradient-based attribution with sensitivity to input changes. The method calculates feature importance by averaging the gradients of the model’s output with respect to the input feature along a path from a reference baseline to the actual input. The baseline is chosen to represent a neutral state, such as a zero vector or a blank image, with no informative features. By aggregating gradients along this path, IG captures how changes in each input feature influence the output, making it applicable to any differentiable model, even when the relationship between inputs and outputs is highly non-linear.
- **Input x gradients (IxGs):** DNNs are highly sensitive to changes made in their inputs. This sensitivity led to the basis of gradient-based attribution methods [136]. One of the simplest and earliest approaches in this category is Input Gradients [137], which explains a model’s prediction by computing the gradient of the output with respect to each input feature. These computed gradients tell the impact of small changes in input features on the model’s output, thereby reflecting the local importance of that feature for a specific prediction. The intuition behind the I*×*G approach is similar to explanations used in linear models, where predictions are interpreted as the product of feature values and their corresponding coefficients. In a comparable way, I×G determines feature attributions by multiplying each input feature value with its local gradient, combining feature magnitude with model sensitivity [138]. Features associated with larger gradient magnitudes are therefore considered to have a stronger impact on the model’s decision. Input gradients are computationally efficient and make use of the internal structure of differentiable models, which makes them well-suited for DNNs.
- **Guided backpropagation (GBP):** GBP [139] is a gradient-based method used to interpret Convolutional Neural Networks (CNNs) by showing which parts of the input most influence the model’s output. It creates saliency maps, which are visual representations highlighting the important regions in the input that contribute strongly to the prediction. GB works by modifying the standard backpropagation algorithm. Normally, gradients are propagated backward through all neurons. In Guided Backpropagation, the algorithm only allows positive gradients to flow [140]. This means that a neuron passes on gradient information only if it was activated positively during the forward pass. Negative gradients are blocked to prevent sending “negative signals” back through the network. The method specifically targets networks that use ReLU activation functions, which are common in CNNs. During the computation of the gradient, GB overrides the usual ReLU gradients so that only non-negative gradients are propagated [141]. This modification helps produce clearer and sharper saliency maps, showing the areas of the input that strongly contribute to the output. Thus, Guided Backpropagation works by highlighting the most relevant input features for a prediction by combining standard gradient information with a filtering step that blocks negative contributions. This makes the resulting visualization easier to interpret and more informative for understanding how the CNN makes decisions.
- **LRP:** LRP [88] is a backpropagation-based method that works by assigning relevance scores to each neuron, showing how much each one contributes to the network’s final output. The process starts at the output layer, where the neuron representing the target class is given full relevance. LRP then works backward through the network layer by layer. At each layer, it distributes the relevance of a neuron to the neurons in the previous layer according to specific rules [142], ensuring that the total relevance is conserved. This iterative process continues until it reaches the input layer. By the time it reaches the first layer, each input neuron has a relevance score that indicates how important that feature was for the final decision. The relevance scores of the input neurons can then be visualized as a heat map, highlighting the features that most strongly influenced the model’s prediction. This makes LRP a powerful tool for interpreting deep networks, as it traces the model’s decision all the way back to the original input features in a way that is easy to understand.
- **Local Interpretable Model-agnostic Explanations (LIME):** Complex ML models often make accurate predictions, but their decision-making process is not easily understandable. LIME is a widely used technique that explains individual model predictions by approximating the complex model locally with a simpler, interpretable surrogate model [85]. Instead of analyzing the entire model, LIME focuses on a specific input instance and studies how small perturbations in its features affect the prediction. By generating variations of the input and observing the corresponding outputs, LIME learns a local linear model whose weights indicate the importance of each feature for that particular prediction. This approach makes LIME model-agnostic, as it does not require access to the internal structure of the model and can be applied to any black-box system. Although LIME provides intuitive and human-understandable explanations, its reliance on sampling and perturbation can introduce instability, and the quality of explanations depends on how well the surrogate model approximates the true model behavior in the local region.
- **SHAP:** SHapley Additive exPlanations (SHAP) [84] comes from Shapley values in cooperative game theory, where a total reward is divided among members of a group according to how much each member contributes. In SHAP, the prediction problem is defined as a game for a single instance of the dataset. The players are the feature values of that instance that work together to produce a prediction. The gain is the model’s prediction for that instance minus the average prediction over all instances. Shapley values measure the average contribution of each feature to this gain by looking at what changes in the prediction when the feature is added to different subsets of other features. To compute this, SHAP evaluates all possible combinations of features, covering cases where all features are present as well as cases where only a subset is used. This makes the method theoretically strong, but the computation becomes very expensive as the number of features increase. To hanDLe this issue, several SHAP variants have been proposed, such as Kernel SHAP, TreeSHAP, and DeepSHAP, which approximate the same idea while reducing the computational cost.
- **DeepLIFT:** DL Important FeaTures (DeepLIFT) [86] explains a model’s prediction by showing how much each input feature contributes to the final output. It works in a way similar to backpropagation, but instead of using gradients, it looks at differences from a reference. For a given input, DeepLIFT compares the model’s output with the output obtained from a neutral or baseline input, called the reference. The same comparison is done at the input level, where each feature is compared to its reference value. The method then assigns a contribution score to each input feature, showing how much changing that feature from its reference value affects the prediction. DeepLIFT then uses the chain rule to pass contribution scores backward through the network, so contributions from deeper layers are correctly distributed to earlier layers. DeepLIFT uses simple rules to do this: a linear rule for layers like fully connected and convolution layers, and a rescale rule for non-linear layers such as ReLU or sigmoid. Overall, DeepLIFT provides a clear local explanation for a single prediction by breaking it into understandable feature-level contributions.
- **GradientSHAP:** GradientShap explains a model’s prediction by showing how much each input feature contributes to the final output. It is based on expected gradients [143], which is an extension of IGs. IGs calculates the contribution of each feature by comparing the model’s output at the actual input to the output at a baseline or reference input [87]. However, it uses only one baseline, which can sometimes limit how well it captures feature effects [144].

GradientShap improves on this by using many baselines sampled from a background dataset and adding small random noise to the inputs. Instead of integrating along a single straight path from baseline to input, it picks a random point along this path multiple times. Then, it calculates the gradient of the model output with respect to the input at these points. By repeating this process many times and averaging, GradientShap produces expected gradients that approximate the Shapley values from game theory. The result is a set of feature attributions that show how much each feature contributes to the difference between the model’s expected output and the actual output for a specific input. One important assumption in GradientShap is that features are independent, which makes calculations simpler but may ignore interactions between features [145]. Thus, GradientShap combines the ideas of IGs and Shapley values. It measures how each feature affects the prediction using gradients and then averages these effects over multiple reference points and noise samples. The result is a set of feature contributions that add up to roughly the difference between the model’s expected output and the actual output, making the explanation clear and easy to understand.

- **DeepSHAP:** DeepSHAP [84] is an advanced feature attribution method that combines ideas from SHAP and DeepLIFT to efficiently explain predictions of DL models. Similar to SHAP, it is based on cooperative game theory and assigns importance scores to input features according to their contribution to the final prediction. However, computing exact Shapley values using traditional SHAP can be computationally expensive for DNNs. To address this, DeepSHAP incorporates the model-specific backpropagation rules of DeepLIFT to approximate Shapley values in a more efficient manner. Thus, DeepSHAP approximates Shapley values using DeepLIFT’s layer-wise decomposition. Using the internal structure of deep networks, particularly convolutional neural networks (CNNs), it can generate pixel- or voxel-level attributions [146] for high-dimensional inputs such as images or biomedical data. Instead of computing Shapley values exactly, DeepSHAP applies DeepLIFT with a reference or background distribution, often called a mask, which defines a baseline input for comparison. Each layer is treated as a linear function, and Shapley decomposition is applied during backpropagation. Unlike DeepLIFT, which uses a single reference and can produce biased attributions, DeepSHAP integrates over multiple background samples to estimate attributions that sum to the difference between the expected model output on the reference distribution and the current model output. By combining the theoretical foundation of Shapley values with DeepLIFT’s computational efficiency, DeepSHAP provides consistent and reliable explanations while remaining practical for large-scale DL applications.
- **Kernel SHAP:** Kernel SHAP is a model-agnostic XAI approach that uses SHAP values from cooperative game theory to measure the importance of each feature in an individual prediction [84].The prediction in this framework is considered as a result of a cooperative game in which each feature plays the role of a player that plays a part in the final prediction. This technique is called Kernel SHAP because the method uses an approximation of the Shapley values using a locally weighted linear regression model and a kernel weighting function. In contrast to model-specific explanation techniques, Kernel SHAP does not need to know the model nor any of its parameters, and so can be used with both ML and DL models. The algorithm satisfies important theoretical properties, including local accuracy, missingness, and consistency, ensuring that the computed feature attributions are mathematically well-defined and comparable across different predictive models.
- **Kernel Explainer:** The Kernel Explainer is the implementation of the Kernel SHAP algorithm offered in the SHAP library, and uses the same theoretical approach as the Kernel SHAP algorithm to estimate feature attributions for black-box predictive models [84]. It creates several perturbed samples with varying combinations of input features, just tests the predictive model on the perturbed samples, and fits a weighted linear regression model to approximate the Shapley value for each feature. The Shapley values that emerge quantify the magnitude and direction of the contributions of each feature, relative to a baseline prediction: positive values indicate that the feature increases the prediction, while negative values indicate a decrease. Beyond the official SHAP implementation, there are several alternative softwares, including the kernelshap package, which share the same theoretical basis but have different sampling and optimization approaches to make computation more efficient [147, 148]. While they can use slightly different ways of computing feature attribution estimates, these implementations follow the same SHAP methodology and offer a consistent and model-agnostic approach to interpreting feature attribution estimates of a predictive model.
- **Permutation Explainer:** Permutation Explainer is another model-agnostic explainer in the SHAP framework that computes feature attributions based on the permutation approach, instead of kernel-weighted regression. The approach is based on full-feature permutations in which the input features are ordered in various sequences and the resulting model predictions are compared to estimate the contribution of each feature to the model prediction. Permutation Explainer lets you directly estimate feature contributions by evaluating your model on a sequence of perturbed samples, unlike Kernel Explainer, which uses a locally weighted linear regression model to approximate Shapley values from perturbed samples. This estimation strategy provides an efficient approximation of Shapley values by leveraging sequential feature permutations to quantify individual feature contributions [84, 149].

Permutation Explainer estimates feature attributions by evaluating the predictive model over multiple forward and reverse permutations of the input features. The contribution of a feature for each permutation is defined as the difference between the model prediction with and without the feature added to the current subset of features. The contributions for each of the permutations that were evaluated are then averaged to estimate the corresponding Shapley value. To be more efficient in computation, the explainer uses intermediate model evaluations when possible, thus decreasing the number of model inferences that must be repeated during the estimation process [150]. Permutation Explainer provides reliable local feature attributions that are consistent with the original SHAP methodology and is efficient in evaluating the model.

- **Feature Permutation:** Feature Permutation is a model-agnostic explainability technique that measures the importance of each input feature based on the change in model performance after randomly permuting its values. The permutation feature importance introduced by Breiman (2001) for random forests [107]. Later, Fisher, Rudin, and Dominici, extended this idea to a general permutation feature importance method applicable to any predictive model [151]. The underlying principle is that if a feature is important for prediction, randomly shuffling its values breaks the relationship between that feature and the target variable, and leads to decrease model performance. Feature permutation first evaluates the trained model on test dataset to establish a baseline performance. The values of one feature are then randomly shuffled while all other features remain unchanged, and the model is evaluated again. The decrease in performance is used as the importance score for that feature. This process is repeated for each feature, and features causing larger performance degradation are considered more influential for the model’s predictions. Feature permutation was developed to overcome the limitations of model-specific feature importance methods that are restricted to particular learning algorithms, such as decision tree impurity-based importance, and gradient-based attribution methods that require access to model gradients and internal parameters. Instead, it directly measures feature importance by evaluating the decrease in model performance after permuting each feature. It does not require feature normalization, can be applied to almost any ML or DL model [152]. However, it may underestimate the importance of highly correlated features, because correlated features can compensate for one another when one is permuted.
- **Saliency:** Saliency [153] is a gradient-based explainability technique that identifies the most influential input features that contribute to output for a specific class. It computes the gradient of the predicted output with respect to the input using backpropagation. These gradients indicate how small changes in each input feature would affect the prediction. They also verify that the model is focusing on relevant information rather than irrelevant patterns or background noise. Since saliency relies on gradient information, it is applicable only to differentiable DL models. Saliency maps are widely used to analyze and improve DL models by showing which input features influence predictions. However, they may produce noisy or unstable explanations and sometimes highlight irrelevant features. In addition, saliency maps indicate which features are important but do not explain why they are important. Therefore, they are often used together with other XAI methods, such as LIME or SHAP, to provide more reliable and comprehensive interpretations.
- **Shapley Value Sampling:** Shapley Value Sampling [154] is an explainability method based on Shapley values from game theory. It tells how much the different features contributed to a specific prediction of a model. Shapley Value Sampling estimates Shapley values by randomly sampling different orders (permutations) of input features instead of evaluating every possible feature combination. For each sampled permutation, it measures how much adding a feature changes the prediction. The average contribution of a feature across many sampled permutations is taken as its Shapley value. Instead of evaluating all possible feature combinations, Shapley Value Sampling estimates feature contributions by randomly sampling different feature orders. This reduces the computational cost but provides an approximate, rather than exact, estimate of the Shapley values.

### 1.6 Evaluation Metrics

- ***Classification:*** Following the evaluation criteria of existing CVDs classification studies, we assess 35 AI classifiers using four evaluation metrics, macro-averaged accuracy (MACC), precision (PR), recall (RC), and F1-score (F1) [155, 156]. Each metric considers all classes equally by computing the measure independently for each class and then averaging the results, thereby ensuring equal contribution from all CVDs classes irrespective of their sample sizes. MACC measures the proportion of correct predictions, both positive and negative, for each class and is computed as the mean accuracy across all individual CVDs classes. Macro PR is the mean of the PR scores across all CVDs classes and is calculated as the ratio of true positives to the total number of instances predicted for each class. Macro RC is calculated as the ratio of true positives to the total number of actual instances of each class and represents the mean RC across all classes. Macro F1-score is obtained by first computing the F1-score for each individual class and then taking the mean of these scores. For each CVDs class, the F1-score is calculated as the harmonic mean of its corresponding PR and RC.

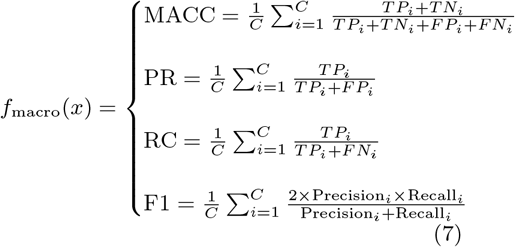 Here, *C* denotes the total number of classes, while *TP*_*i*_, *TN*_*i*_, *FP*_*i*_, and *FN*_*i*_ represent the true positives, true negatives, false positives, and false negatives for the *i*-th class, respectively.
- ***XAI:*** The quality and robustness of the explanations are evaluated using 3 distinct XAI evaluation metrics namely, area over the perturbation curve (AOPC) [89], infidelity [90], and maximum sensitivity [90]. AOPC [89] measures faithfulness of an explanation to the model’s prediction. It measures how quickly the model’s output changes when input features are perturbed or removed in a specific order determined by their assigned importance (either most relevant first (MoRF) or least relevant first (LeRF)). Infidelity [90] measures how well an explanation *ϕ* captures the change in a model’s prediction when significant perturbations are applied to the input. Formally, infidelity is defined as the expected mean squared error between the change predicted by the explanation and the actual change in the model output under significant perturbations of the input, thereby measuring the unfaithfulness of the explanation to the model’s true behavior. Sensitivity [90] refers to the stability of an explanation, when insignificant perturbations are applied to the input. It measures the maximum degree to which an explanation is affected when the input is slightly modified.

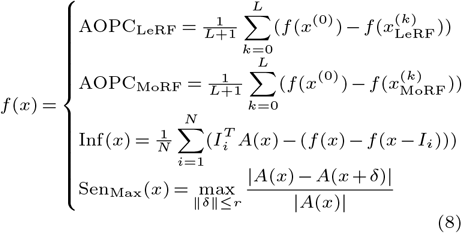 In AOPC, where *f* (*x*^(0)^) is the prediction score of the original input, 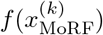 and 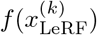 are the prediction scores after removing the *k* most relevant and least relevant features, respectively, and *L* denotes the total number of perturbation steps. In the infidelity, *A*(*x*) represents the attribution vector, *I* shows the perturbation vector, *f* (*x*) denotes the model prediciton for the original input, *f* (*x* − *I*) represents the model prediciton for the perturbed input, and (*f* (*x*)−*f* (*x*−*I*_*i*_)) shows the actual output change. In the sensitivity max metric, *A*(*x*) represent the attributions of the original input, *A*(*x* + *δ*) denotes post perturbation attributions, *r* shows the perturbation radius.
- ***Watermelon Multisection Evaluation:*** Harmonization quality was interpreted using rrrr3the ratio 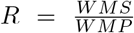, where *WMS* denotes the WM metric calculated with respect to biological sample classes and *WMP* denotes the metric calculated with respect to experimental platform classes. Since effective harmonization should reinforce biological similarity while reducing platform-driven separation, higher values of *R* indicate better harmonization performance. By comparing tissue-based and platform-based WM metrics, this approach helps determine which harmonization technique most effectively minimizes batch effects while preserving biologically meaningful variation.

## 2 Results

### 2.1 Harmonization of Transcriptomic Datasets

Figures 2A–C present the harmonization performance of Shambhala-2, TDM, and UPC across RNA-seq datasets from blood, left ventricle, and heart tissues, with the WM ratio used to assess each method’s ability to minimize platform-driven variation while preserving biologically meaningful clustering. Figure 2A illustrates the results for blood datasets, where Shambhala-2 achieves the highest median WM ratio (1.714), outperforming TDM (1.333) and UPC (1.286), indicating superior preservation of biological structure following harmonization. Figure 2B presents the left ventricle datasets and shows a similar trend, with Shambhala-2 again producing the highest median WM ratio (1.667), whereas TDM and UPC yield lower values of 1.250 and 1.000, respectively. Figure 2C displays the results for heart datasets, where performance differences among the methods are less pronounced; however, Shambhala-2 maintains the highest median WM ratio (1.333), followed by TDM (1.243) and UPC (1.214). Overall, Shambhala-2 consistently achieves the highest WM ratios across all RNA-seq tissue groups, suggesting more effective reduction of platform-related variation while preserving biologically relevant clustering, whereas TDM demonstrates intermediate performance and UPC exhibits greater variability and generally lower effectiveness.

**Figure 2:**
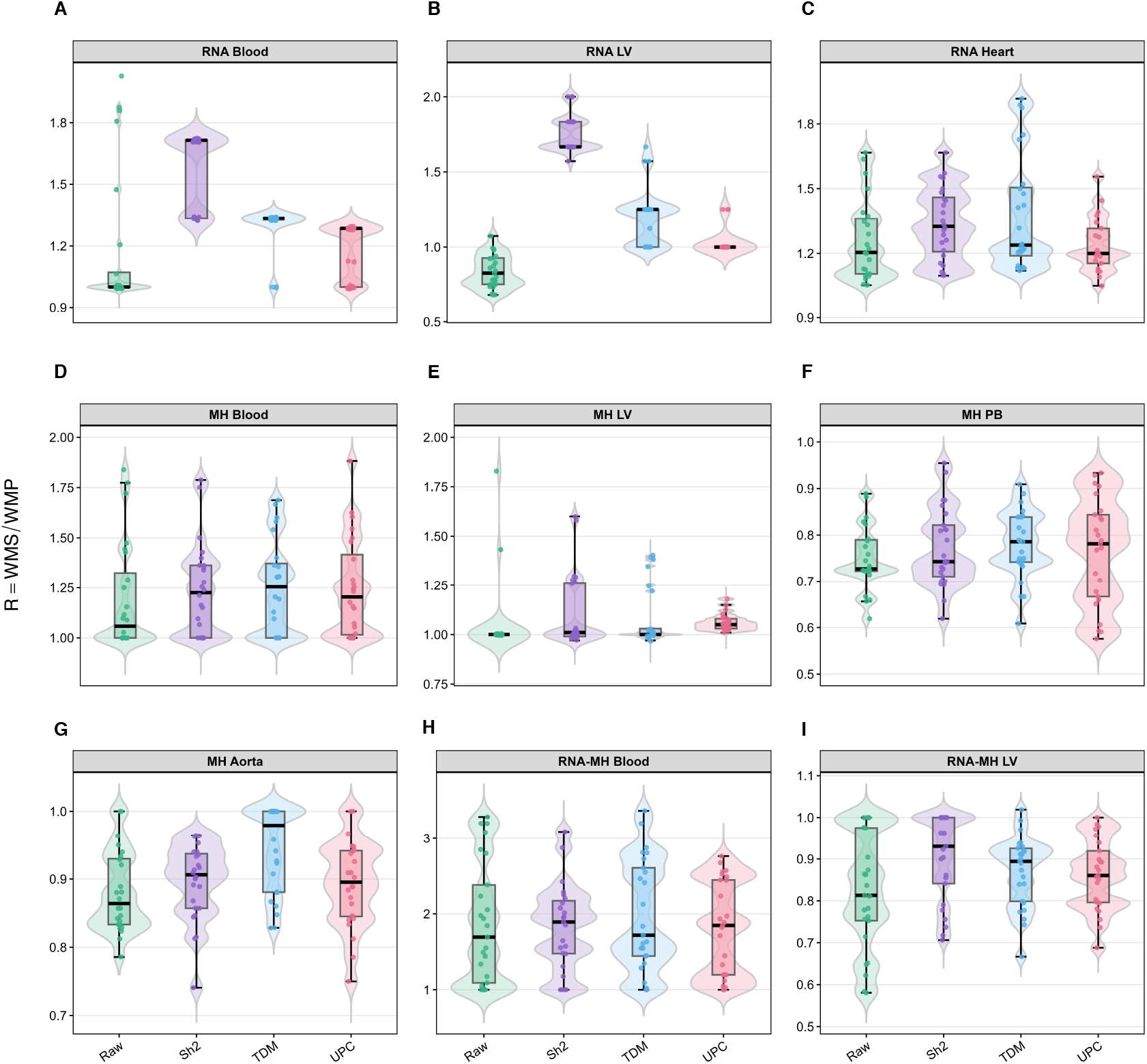
Watermelon evaluation across all normalization methods and datasets. Violin plots represent the distribution of WM ratios for Raw, Shambhala2, TDM, and UPC normalized data across RNA-seq and MH tissues. Boxplots indicate the median and interquartile range, while individual points represent repeated WM evaluation iterations. Higher WM ratios indicate stronger preservation of biological signal relative to platform-associated batch effects.

Figures 2D–G present the WM ratio of Shambhala-2, TDM, and UPC across microarray hybridization (MH) datasets from blood, left ventricle, peripheral blood, and aorta tissues. Figure 2D illustrates the results for blood datasets, where all three methods achieve median WM ratios greater than 1, indicating that biological signal remains stronger than platform-driven variation following harmonization. Shambhala-2 produces the highest median WM ratio (1.227), followed by TDM (1.194) and UPC (1.161), although the differences among methods are relatively small. Figure 2E shows the harmonization performance for left ventricle datasets, where WM ratios are close to 1.0 for all methods, with UPC achieving the highest median value (1.050), followed by Shambhala-2 (1.010) and TDM (1.000), which suggests a comparable contribution of biological and platform-related factors to the observed clustering patterns. Figure 2F presents the results for peripheral blood datasets, where all methods yield median WM ratios below 1, indicating that platform effects remain more prominent than biological signal after harmonization. TDM achieves the highest median WM ratio (0.786), followed closely by UPC (0.781) and Shambhala-2 (0.743). Figure 2G displays the harmonization performance for aorta datasets, where median WM ratios also remain below 1 but are relatively similar across methods, with TDM achieving the highest value (0.958), followed by Shambhala-2 (0.903) and UPC (0.889). Overall, the three harmonization methods exhibit more comparable performance across MH datasets than in RNA-seq datasets, with modest differences in WM ratios across tissues, suggesting that platform-related variation is less strongly affected by the choice of harmonization method in microarray data.

Figures 2H–I present the harmonization performance of Shambhala-2, TDM, and UPC on the combined RNA-seq and microarray datasets from blood and left ventricle tissues. Figure 2H illustrates the results for blood datasets, where all methods achieve WM ratios substantially greater than 1.0. Shambhala-2 yields the highest median WM ratio (1.95), followed by UPC (1.84) and TDM (1.72), suggesting slightly stronger preservation of biological structure during cross-platform integration. Figure 2I shows the harmonization performance for left ventricle datasets, where WM ratios remain close to 1.0, indicating limited separation between biological and platform effects. Shambhala-2 again achieves the highest median WM ratio (0.93), followed by TDM (0.89) and UPC (0.86), although the substantial overlap among the distributions suggests broaDLy comparable performance among the three methods. Overall, Shambhala-2 consistently achieves the highest median WM ratios in both cross-platform tissue groups, which indicates a modest advantage in preserving biological signal while reducing platform-related variation during integration of RNA-seq and microarray datasets.

Overall, Shambhala-2 achieves the highest median WM ratios across all RNA-seq tissue groups and maintains competitive performance across microarray datasets. In addition, it consistently yields the highest WM ratios in the cross-platform blood and left ventricle datasets which shows effective reduction of platform-related variation while preserving biologically meaningful clustering. These results suggest that Shambhala-2 provides the most reliable harmonization of heterogeneous transcriptomic datasets and is therefore the most appropriate method for downstream analyses.

### 2.2 Classification Performance of ML and DL Classifiers

Fig. 3a illustrates the mean F1-scores achieved by 15 distinct ML classifiers across the MH, RNA, and combined RNA/MH modalities using the Shambhala-harmonized datasets. In the MH modality, HGB achieves the highest F1-score (0.777), followed by LR (0.683) and RF (0.667), indicating strong predictive performance. Similar results are observed for the RNA modality, where HGB again achieves the highest F1-score (0.777), with LR (0.739), RF (0.738), and SVM (0.735) performing comparably well. For the combined RNA/MH modality, XGB achieves the highest F1-score (0.724), followed by LR (0.713) and HGB (0.697). Across all modalities, ensemble-based methods, including HGB, RF, and XGB, together with linear models such as LR and SVM, consistently rank among the best-performing classifiers. In contrast, probabilistic classifiers and several boosting-based approaches generally achieve lower F1-scores, particularly in the combined modality. These results suggest that ensemble and linear methods provide the most robust classification performance on harmonized transcriptomic datasets.

**Figure 3:**
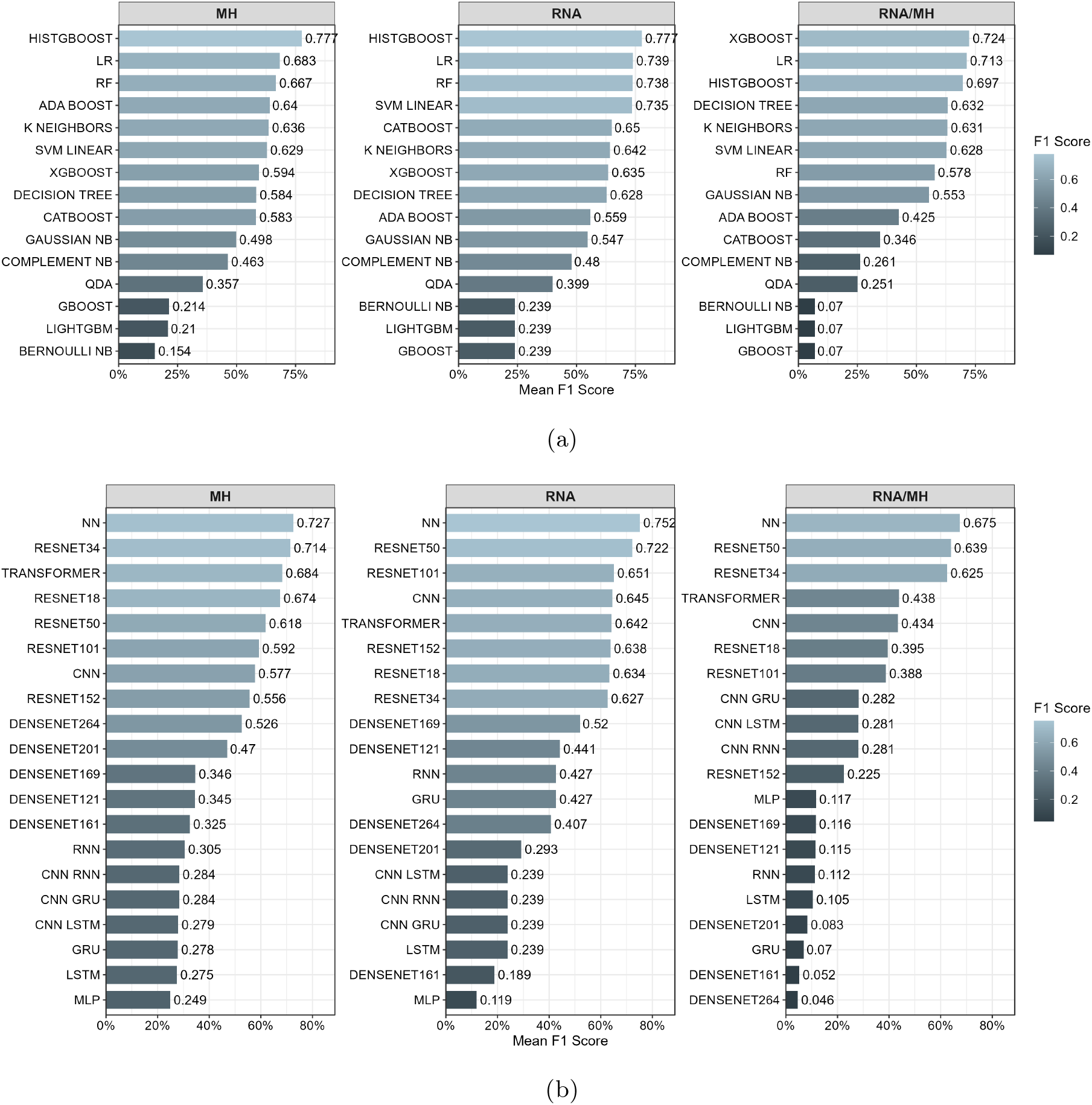
(a) Performance of ML classifiers across datasets of MH, RNA and RNA/MH modalities based on macro-average F1-score. (b) Performance of DL models across the same datasets. The results provide a comparative view of model effectiveness across datasets, highlighting variations in performance across classifiers and data types.

Fig. 3b presents the mean F1-scores achieved by 20 distinct DL classifiers across the MH, RNA, and combined RNA/MH modalities using the Shambhala-harmonized datasets. In the MH modality, NN achieves the highest F1-score (0.727), followed by ResNet34 (0.714) and Transformer (0.684), while other ResNet variants also demonstrate competitive performance. A similar trend is observed in the RNA modality, where NN again achieves the highest F1-score (0.752), followed by ResNet50 (0.722), ResNet101 (0.651), CNN (0.645), and Transformer (0.642). For the combined RNA/MH modality, NN maintains the best performance (0.675), along with ResNet50 (0.639) and ResNet34 (0.625). Across all modalities, feedforward NN and ResNet-based architectures consistently achieve the highest F1-scores, indicating strong generalization and robustness for disease classification. In contrast, DenseNet, recurrent-based, and hybrid architectures generally exhibit lower performance, with the largest declines observed in the combined RNA/MH modality, suggesting greater difficulty in learning discriminative representations from integrated multi-platform data.

### 2.3 XAI results

Figure 4 illustrates AOPC scores for 12 XAI methods applied to the top three DL models (Transformer, NN, and ResNet34) across the RNA, MH, and combined RNA/MH datasets. Shapley Value Sampling ranks highest in nearly every tissue and modality: its MoRF curves rise sharply and separate clearly from the corresponding LeRF curves in the RNA blood, heart, and left ventricle datasets, and this pattern repeats across the MH aorta, blood, left ventricle, and peripheral blood tissues regardless of the underlying model. Input *×* Gradient and IG follow as a second tier, tracking Shapley Value Sampling’s separation pattern but at consistently lower AOPC magnitudes, while LIME, LRP, Feature Permutation, and DeepLIFT trail behind with narrower MoRF-LeRF gaps, pointing to weaker recovery of the truly influential features. The ranking among methods stays stable even as the absolute AOPC values shift across tissues, and the gap between the leading and trailing methods widens further in the combined RNA/MH setting, where Shapley Value Sampling, Input *×* Gradient, and IG remain ahead most clearly in the blood and left ventricle datasets. Taken together, these results position Shapley-based sampling as the most dependable XAI method for the DL models across all three modalities, with Input *×* Gradient and IG as reliable runners-up.

**Figure 4:**
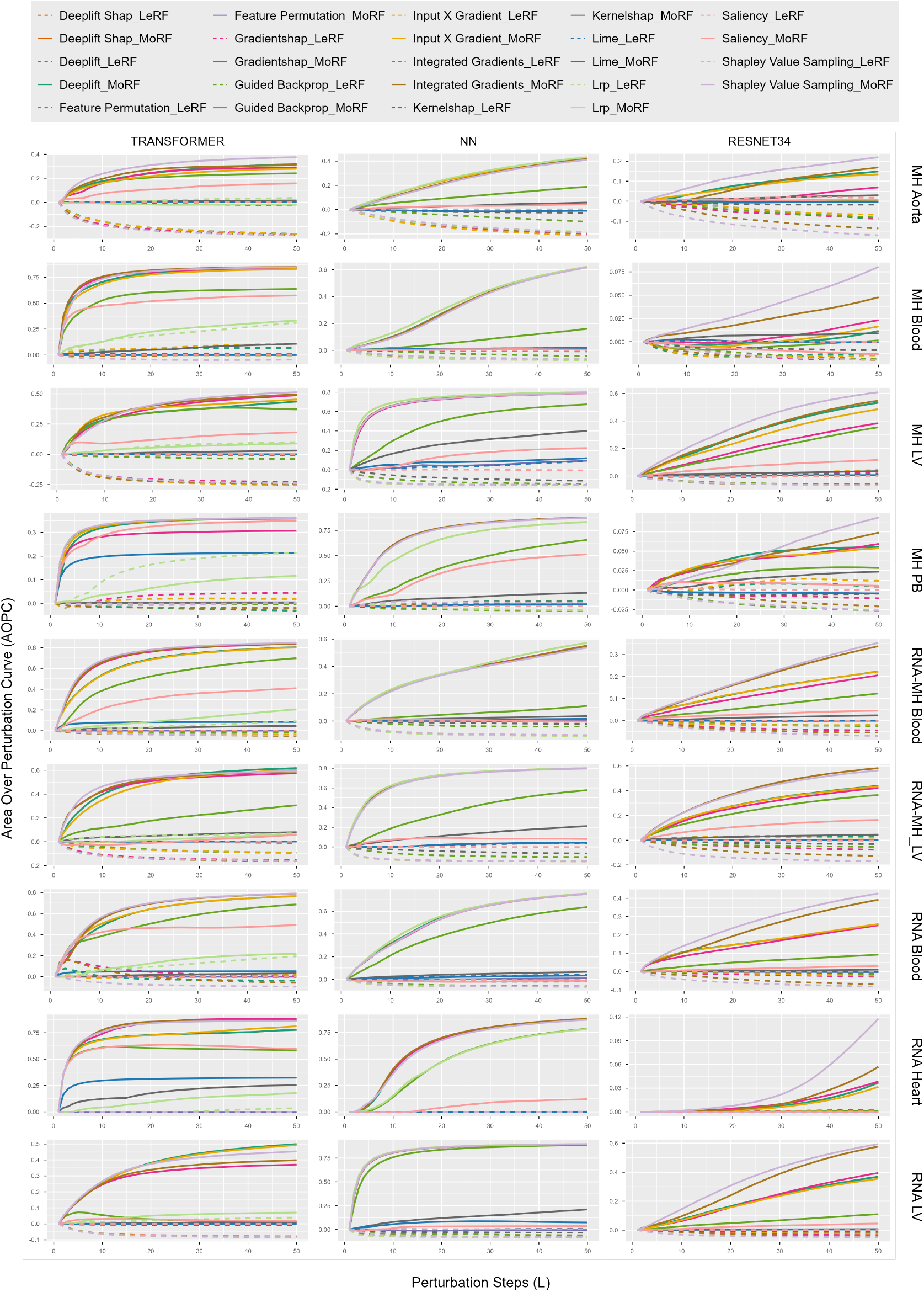
AOPC curves for different datasets (rows) and DL models (columns). Solid lines represent MoRF and dashed lines represent LeRF. Each subplot shows perturbation-based evaluation across 50 perturbation steps.

Figure 5 shows the corresponding AOPC evaluation for 4 XAI methods across the four ML models (HGB, LR, RF, and SVM) and all 9 datasets. Most model–dataset pairs show a clean split between the MoRF and LeRF curves, confirming that the underlying explanations generally track features that matter to the prediction. Permutation Explainer stands out with the widest split, an effect most visible for RF and HGB, indicating that the features it ranks highly are also the ones driving predictive performance; Kernel Explainer reaches similar separation for a number of LR and SVM models, though less reliably across the full dataset set. Kernel SHAP sits at the other end of the spectrum: across many datasets its MoRF and LeRF trajectories stay close together, occasionally almost flat, suggesting its rankings barely move model output under perturbation. This places Kernel SHAP as the least faithful of the three SHAP-family approximations tested here, while the two permutation-based explainers, Permutation Explainer and Kernel Explainer, emerge as the most trustworthy choices for the ML models.

**Figure 5:**
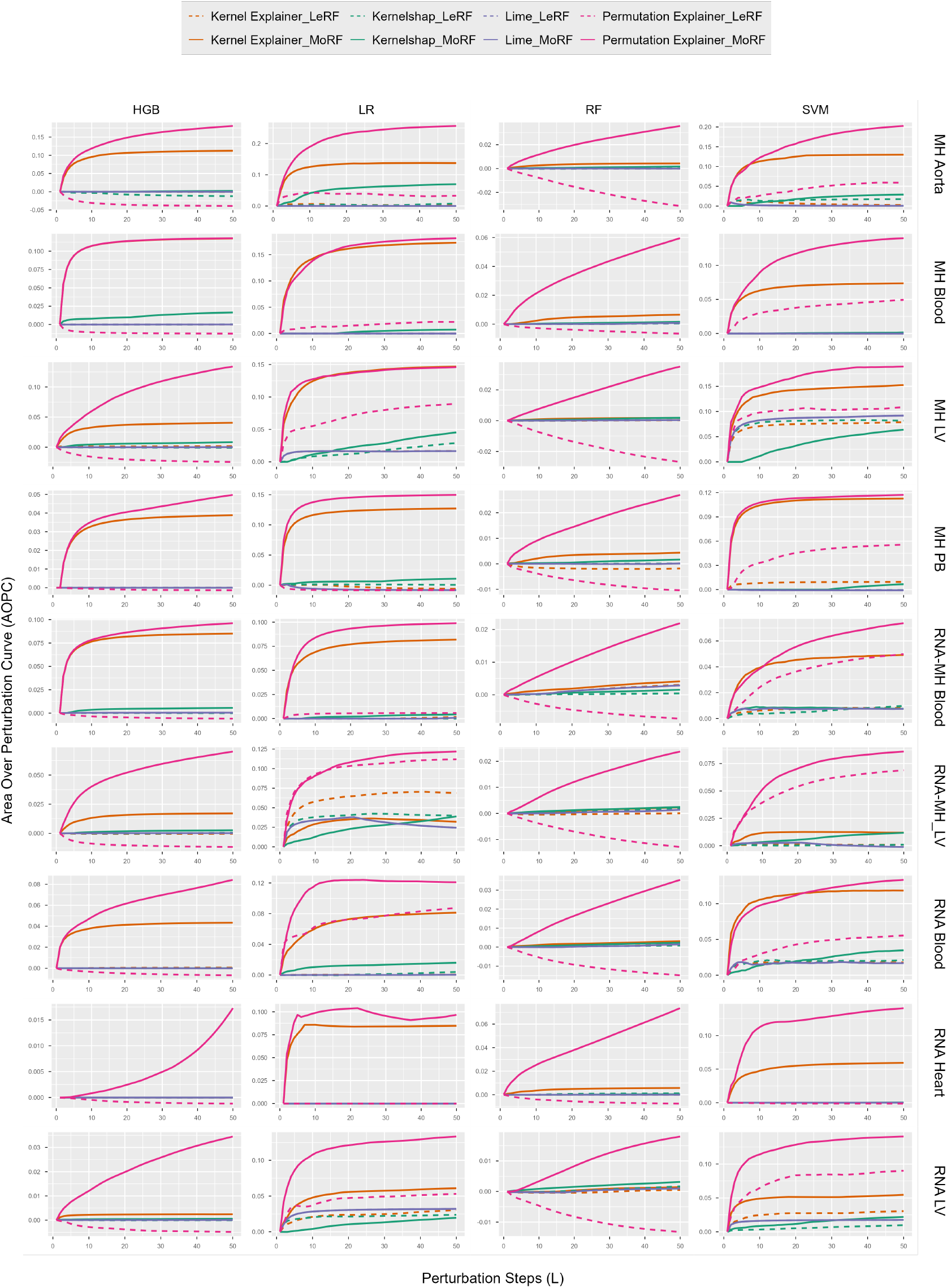
AOPC curves for different datasets (rows) and ML models (columns). Solid lines represent MoRF and dashed lines represent LeRF. Each subplot shows perturbation-based evaluation across 50 perturbation steps.

Figure 6a turns to explanation stability, measured via sensitivity, and exposes a clear architecture effect. NN posts the lowest sensitivity scores, and the tightest distributions, in most datasets (MH Aorta, MH Blood, MH LV, RNA Blood, RNA Heart, RNA LV), meaning its explanations barely move under small input perturbations; this holds for both single-disease and multiclass settings. DeepGene Transformer sits between the other two models, with sensitivity scores above NN but below ResNet34, and its spread widens noticeably in MH PB, RNA-MH Blood, and RNA-MH LV. ResNet34 shows the opposite profile to NN: the highest sensitivity and the broadest distributions of the three, an effect that intensifies further in the combined multiclass datasets. Sensitivity alone therefore favors NN, places DeepGene Transformer in the midDLe, and marks ResNet34 as the least stable of the three.

**Figure 6:**
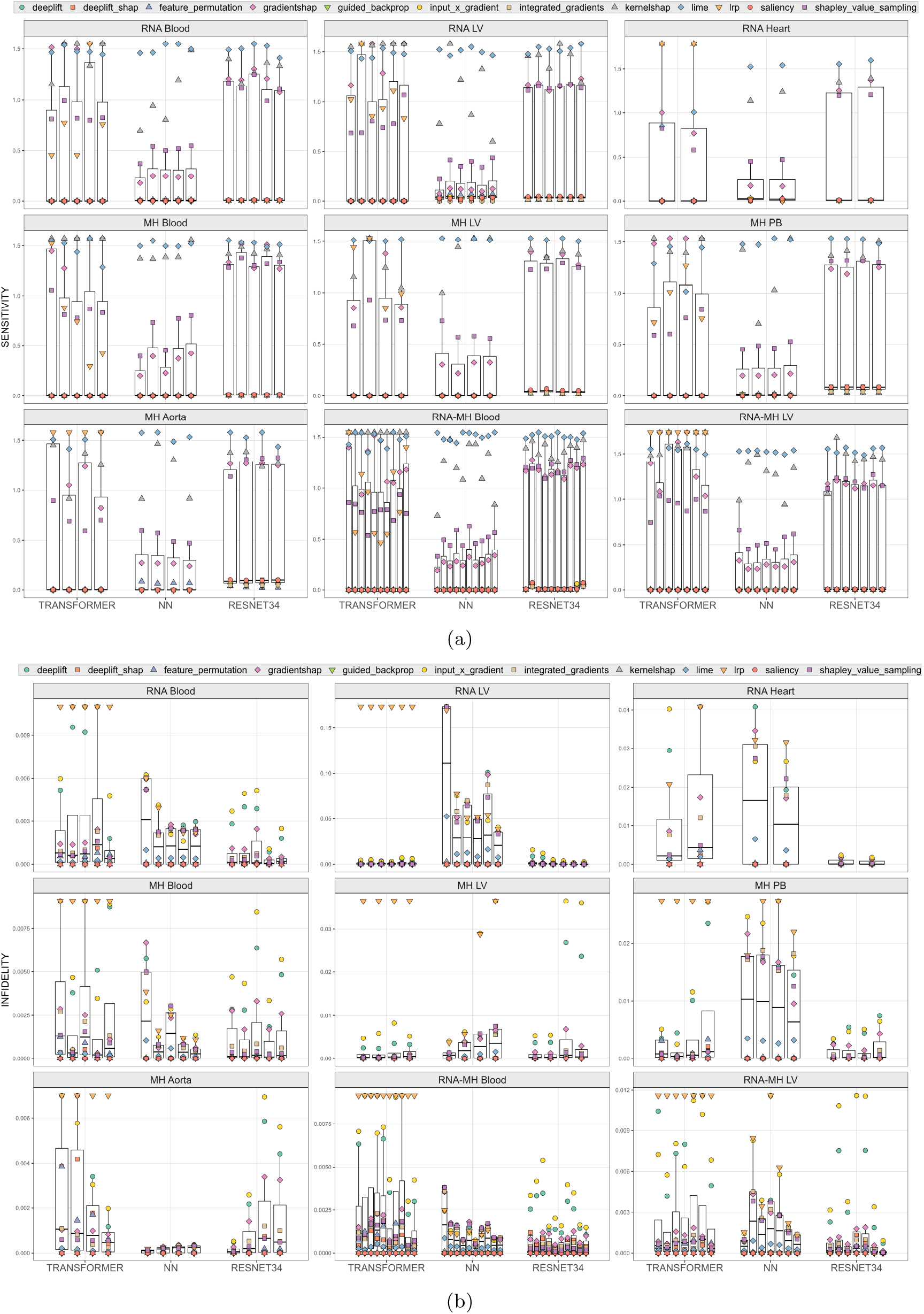
Evaluation of explainability methods across 9 datasets using 3 DL models (Transformer, NN, and ResNet34). (a) Sensitivity and (b) Infidelity results are shown as class-wise grouped boxplots, with colored markers denoting individual methods. Lower values indicate better performance.

Figure 6b evaluates the same three models on infidelity, and the ranking flips relative to sensitivity. ResNet34 now returns the lowest, and most tightly distributed, infidelity scores in most datasets, most clearly in MH Aorta, MH Blood, RNA Blood, RNA Heart, and RNA-MH LV, indicating explanations that better track the model’s actual behavior. DeepGene Transformer again lands close behind, with its median infidelity near ResNet34’s aside from added spread in MH PB and RNA Heart. NN, by contrast, produces the highest infidelity of the three, most sharply in RNA LV and, to a lesser extent, MH PB and RNA Heart, meaning that the stability it showed in Fig. 6a comes at the cost of faithfulness here. Infidelity therefore favors ResNet34, keeps DeepGene Transformer in second place, and leaves NN as the least faithful of the three despite its earlier stability advantage.

Together, the AOPC, sensitivity, and infidelity results reveal a stability–faithfulness trade-off among the DL models that no single architecture wins outright, and they establish Shapley-based sampling and permutation-based explainers as the most reliable attribution choices for the DL and ML models, respectively.

### 2.4 Identified Biomarkers across CVDs

To identify robust biomarkers, the best-performing ML and DL models and their corresponding XAI methods are selected based on AOPC across all datasets. Figure 7 shows the selected models together with their corresponding XAI methods. The selected ML models include RF, SVM, LR, and HistGradientBoost, explained using Permutation and Kernel Explainers. The selected DL models include NN, ResNet34, and Transformer, and their corresponding selected XAI methods are IxG, IG, DeepLIFT, GradientSHAP, and SHAP.

**Figure 7:**
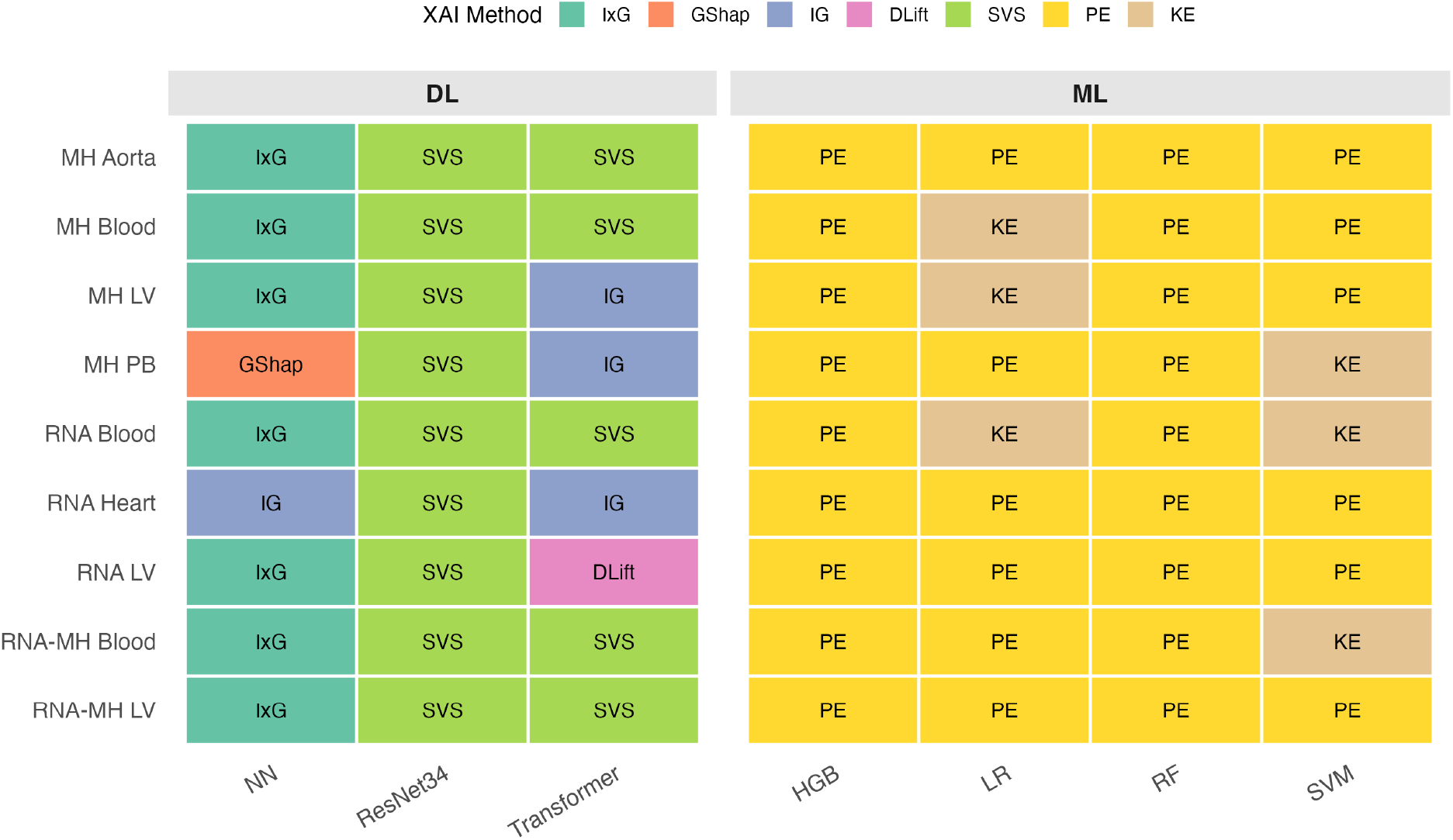
XAI method applied to each dataset–model combination. Each tile shows the selected XAI method used to interpret a given model’s predictions on a given dataset, split into DL models (NN, ResNet34, Transformer) on the left and classical ML models (HGB, LR, RF, SVM) on the right.

For each CVD, the top 10 positive and top 10 negative biomarkers are extracted separately from every selected model–XAI combination, across all datasets. Genes recurring across this collection of datasets, models, and XAI methods are pooled into a consensus list and scored using the procedure summarized in Algorithm 1. For each candidate gene, the algorithm computes: gene frequency (*F*), the number of dataset–model–XAI combinations in which the gene appears, reflecting how consistently it is identified across independent analyses; dataset count (*D*), the number of independent datasets supporting the gene across different tissues and platforms, reflecting biological generalizability; model count (*M*), the number of distinct ML/DL architectures identifying the gene, reflecting robustness to model choice; XAI method count (*X*), the number of distinct explanation techniques highlighting the gene, reflecting robustness to the interpretation method; and mean absolute feature importance (*I*), quantifying the gene’s overall contribution to disease prediction regardless of direction. The consensus score is computed as the product of these five terms (Equation 9), such that a high score requires consistency across *all* dimensions rather than strength along any single one.

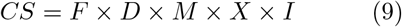

where *CS* denotes the consensus score, *F* the gene frequency, *D* the dataset count, *M* the model count, *X* the XAI method count, and *I* the mean absolute feature importance.

Genes are ranked by consensus score, and the top 20 consensus biomarkers for each CVD are selected separately for the ML and DL analyses. Mean feature importance then determines biomarker direction, classifying each gene as a positive disease biomarker or a negative predictive biomarker. The final consensus biomarkers for all CVDs are listed in Tables 2 and 3 for DL and ML, respectively, with each table organized into three categories. *Disease-associated biomarkers* are genes with consistently positive attribution scores across the selected XAI methods, indicating a positive contribution to disease prediction. *Negative predictive biomarkers* are genes with consistently negative attribution scores; several of these may play protective or compensatory roles, though this requires further biological validation. *Potential biomarkers* are genes additionally supported by published literature or existing biological evidence, representing priority candidates for future experimental and clinical validation.

**Table 2:**
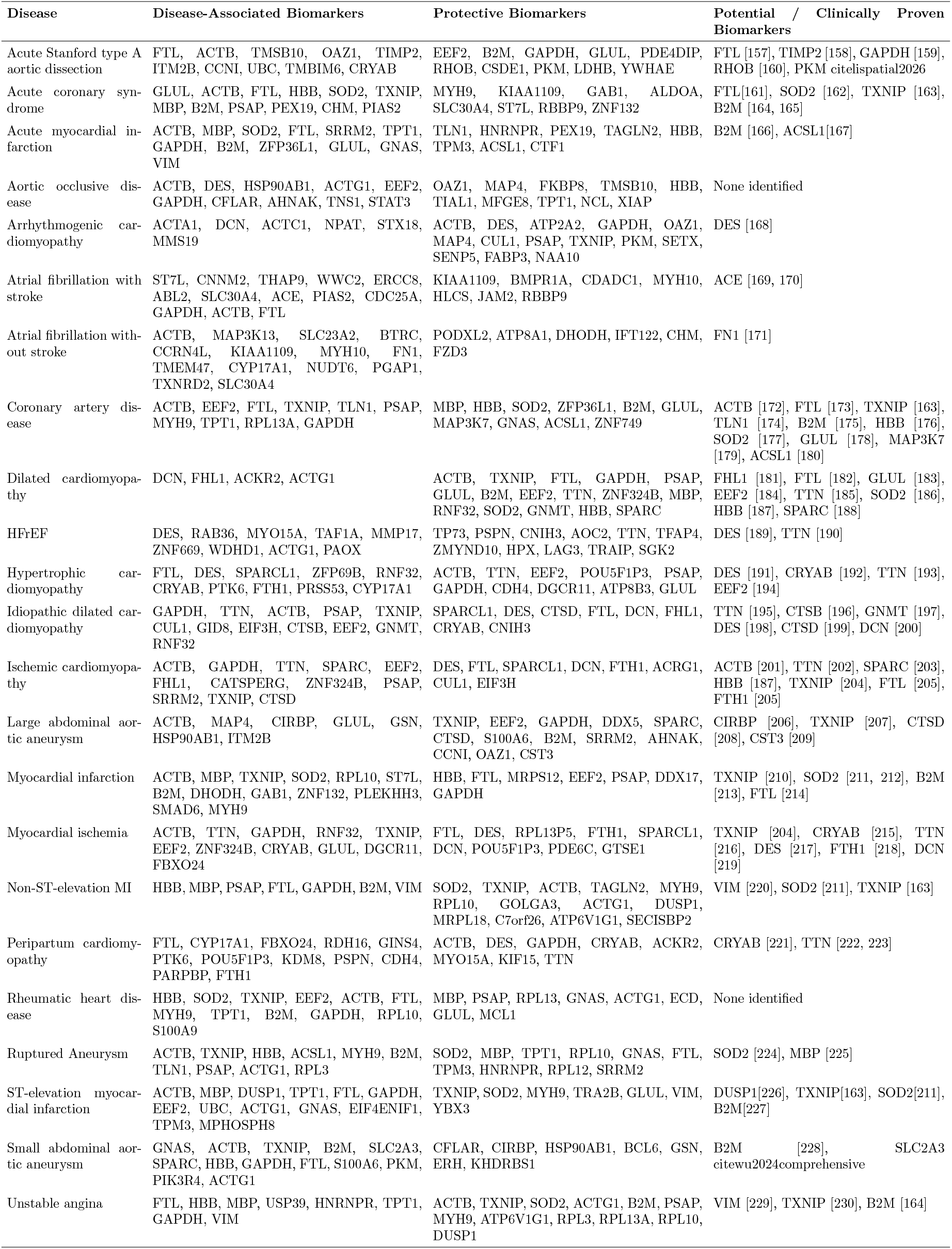
DL-derived biomarkers associated with CVDs.

**Table 3:**
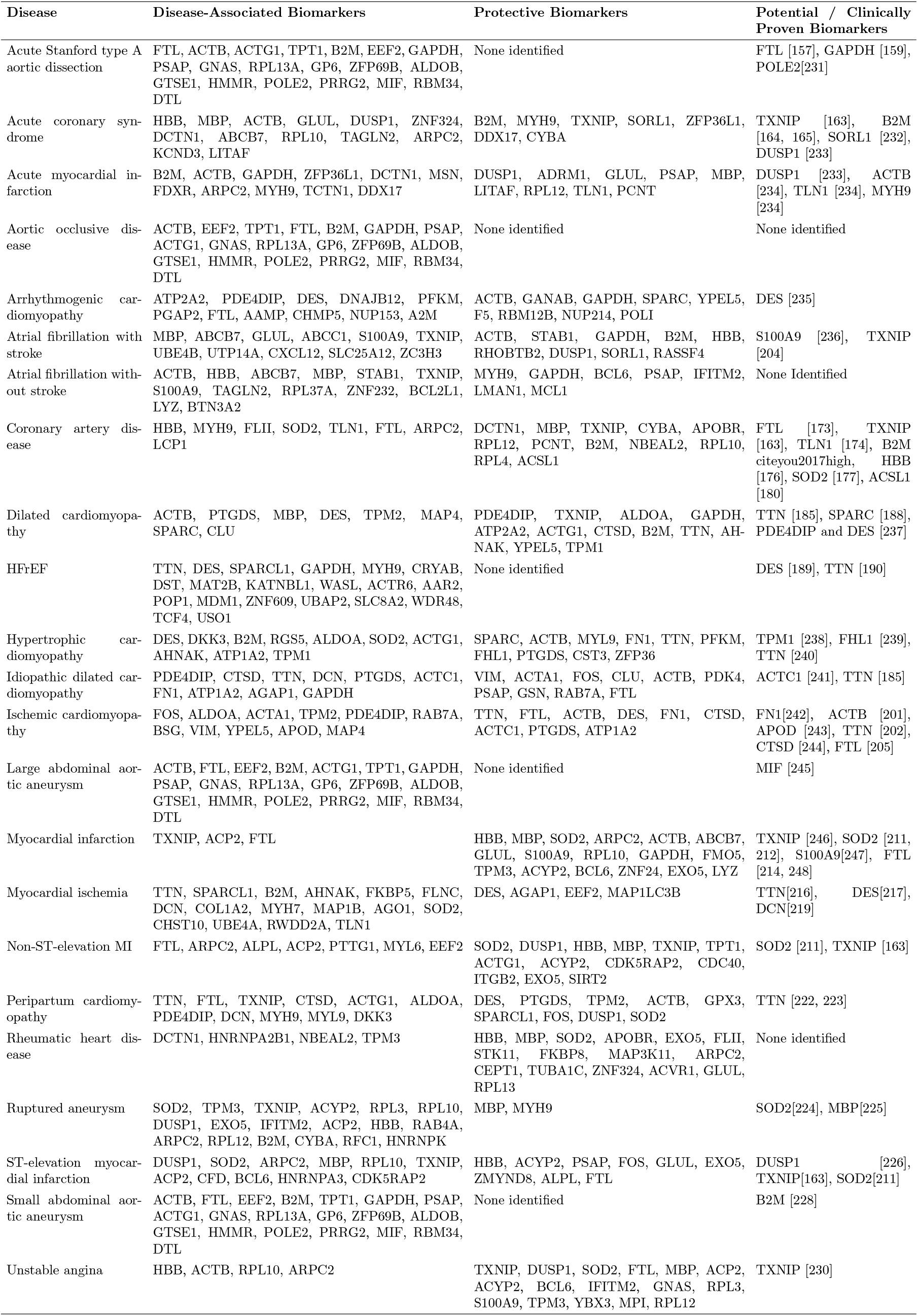
ML-derived biomarkers associated with CVDs.

Across both tables, several genes recur across biologically related disease groups, pointing to shared molecular mechanisms rather than disease-specific artifacts. Among aortic diseases – acute Stanford type A aortic dissection, aortic occlusive disease, ruptured aneurysm, and abdominal aortic aneurysms – FTL, ACTB, TXNIP, SPARC, B2M, and HBB recur most frequently, consistent with shared involvement of inflammation, extracellular matrix remodeling, and cellular stress responses. Ischemic disorders – including acute coronary syndrome, coronary artery disease, myocardial infarction, myocardial ischemia, unstable angina, and ST- and non-ST-elevation myocardial infarction – show frequent enrichment of ACTB, GAPDH, FTL, TXNIP, MBP, HBB, and B2M, genes previously linked to metabolic dysregulation, oxidative stress, and immune activation in ischemic cardiac injury. Among cardiomyopathies, TTN is consistently prioritized in dilated cardiomyopathy, HFrEF, idiopathic dilated cardiomyopathy, and peripartum cardiomyopathy, in line with its established role in myocardial structural integrity and inherited cardiomyopathies, while DES emerges as a prioritized biomarker in arrhythmogenic cardiomyopathy, reflecting the centrality of desmosomal and cytoskeletal abnormalities to that disease’s pathogenesis.

For atrial fibrillation, with and without stroke, and rheumatic heart disease, the prioritized biomarkers instead cluster around inflammatory and immune-related pathways, including ACE, ACTB, FTL, JAM2, MYH10, FN1, and S100A9. Taken together, Tables 2 and 3 reveal substantial heterogeneity in biomarker profiles across CVDs, alongside recurrent genes shared among mechanistically related disease groups. This combination of disease-specific and cross-cutting signals provides a focused set of candidates for future mechanistic studies aimed at improving cardiovascular diagnosis and prognosis.

Among aortic diseases, including acute Stanford type A aortic dissection, aortic occlusive disease, ruptured aneurysm, and abdominal aortic aneurysms, several recurrent genes are observed, particularly FTL, ACTB, TXNIP, SPARC, B2M, and HBB, suggesting common molecular mechanisms related to inflammation, extracellular matrix remodeling, and cellular stress responses. Ischemic disorders, including acute coronary syndrome, coronary artery disease, myocardial infarction, myocardial ischemia, unstable angina, and ST- and non-ST-elevation myocardial infarction, show frequent enrichment of ACTB, GAPDH, FTL, TXNIP, MBP, HBB, and B2M. These genes have previously been associated with metabolic dysregulation, oxidative stress, and immune activation in ischemic cardiac injury. Among cardiomyopathies, TTN was consistently prioritized in dilated cardiomyopathy, HFrEF, idiopathic dilated cardiomyopathy, and peripartum cardiomyopathy, consistent with its established role in myocardial structural integrity and inherited cardiomyopathies. In arrhythmogenic cardiomyopathy, DES emerged as a prioritized biomarker, reflecting the importance of desmosomal and cytoskeletal abnormalities in disease pathogenesis.

#### Algorithm 1

Consensus Score Computation

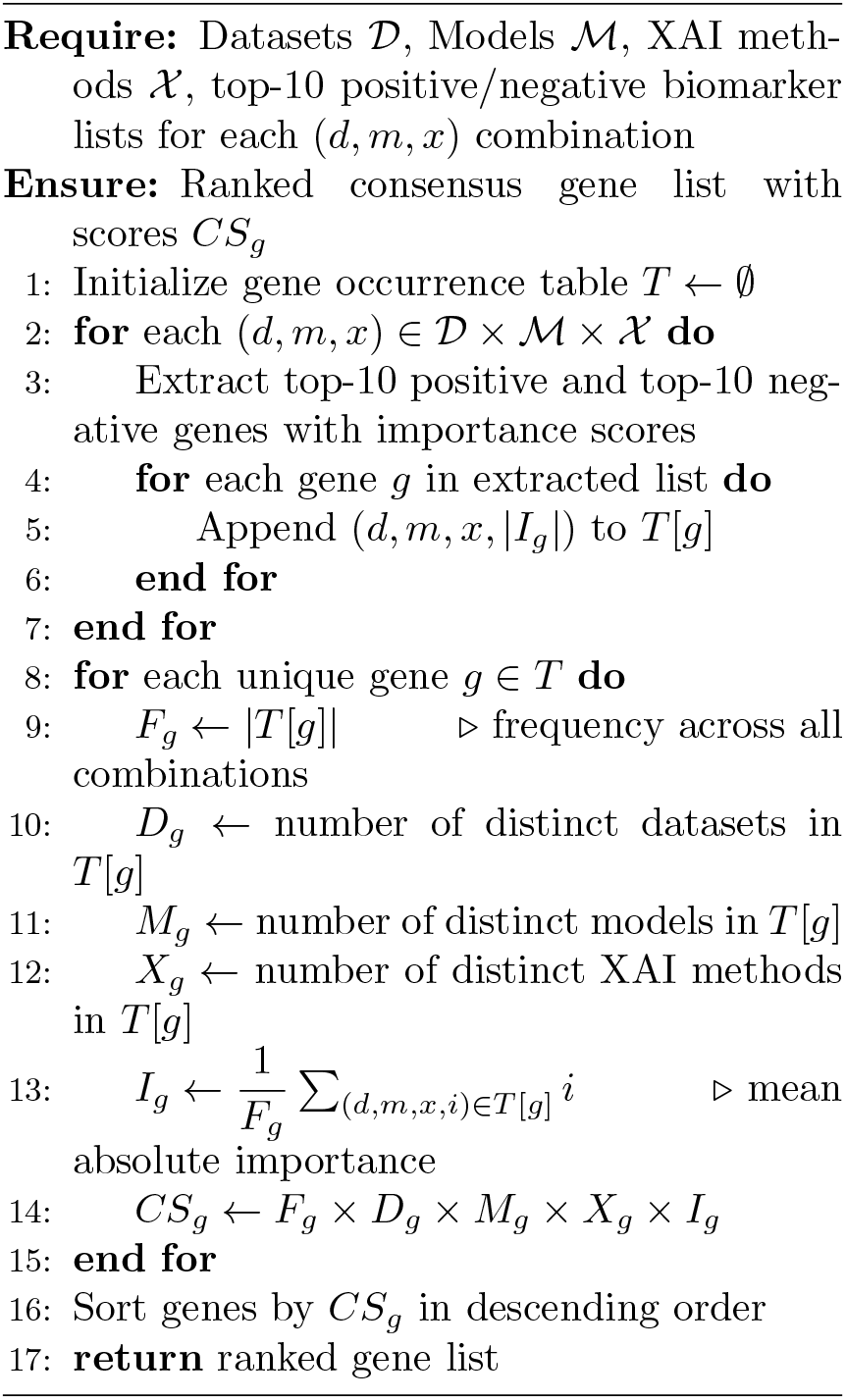

For atrial fibrillation with and without stroke and rheumatic heart disease, the prioritized biomarkers included genes involved in inflammatory and immune-related pathways, such as ACE, ACTB, FTL, JAM2, MYH10, FN1, and S100A9. Overall, the results demonstrate substantial heterogeneity in biomarker profiles across CVDs while also revealing recurrent genes shared among related disease groups. The identified biomarkers provide a focused set of candidates for future mechanistic studies aimed at improving cardiovascular diagnosis and prognosis.

### 2.5 Downstream Analysis

#### Functional Enrichment of Consensus Biomarkers

To assess the biological relevance of the prioritized biomarkers, the consensus gene sets identified by the ML and DL frameworks are separately subjected to Gene Ontology (GO) and KEGG pathway enrichment analysis (Fig. 7). Across both frameworks, the enriched terms converge on cardiac muscle structure, cytoskeletal organization, and cardiomyopathy-related pathways, indicating that the prioritized genes are mechanistically relevant to cardiac dysfunction. At the same time, the two gene sets diverge in several respects, pointing to complementary biological signals captured by the ML and DL models.

#### GO Biological Process

For the ML-derived biomarkers (Fig. 7a), the most enriched terms are muscle contraction, muscle system process, and actin filament organization, followed by muscle organ development and regulation of actin filament-based process. Several of the genes underlying this enrichment, including TTN and DES are also independently prioritized as biomarkers of dilated cardiomyopathy, arrhythmogenic cardiomyopathy, and related structural cardiomyopathies in Table 2, which supports a shared contractile-dysfunction mechanism across these conditions. The ML set additionally enriches endothelial cell migration and response to oxidative stress/reactive oxygen species, processes plausibly linked to the aortic and ischemic disease groups in the cohort, where vascular wall injury and oxidative stress are established contributors to pathology. The DL-derived biomarkers (Fig. 7b) similarly enrich muscle contraction, muscle system process, and muscle organ development, but additionally identify platelet activation, platelet aggregation, homotypic cell–cell adhesion, and wound healing processes centered on genes such as ACTB and FN1, which Table 2 links to atrial fibrillation and myocardial infarction, conditions in which platelet-driven thrombosis and vascular repair are mechanistically central. This suggests that the DL framework captures thrombotic and reparative biology that is less prominent in the ML-derived set.

#### GO Molecular Function

Both gene sets enrich actin binding, actin filament binding, and structural constituent of cytoskeleton (Fig. 7a–b, panel c), consistent with the contractile-apparatus genes (e.g., TTN, DES, ACTG1) shared across the cardiomyopathy classes in Table 2. The ML set additionally enriches structural constituent of muscle, indicating a somewhat stronger emphasis on muscle integrity, whereas the DL set shows comparatively higher p.adjust values for actin binding despite a similar gene count, suggesting a slightly weaker statistical signal for this term in the DL-derived set.

#### GO Cellular Component

Focal adhesion and cell-substrate junction are the most enriched components in both gene sets, followed by myofibril and contractile muscle fiber, again reflecting the myocardial structural genes shared across cardiomyopathy and ischemic phenotypes. The ML set additionally enriches striated muscle thin filament, actin filament, and myofilament, while the DL set uniquely enriches intercalated disc, a structure specifically relevant to arrhythmogenic cardiomyopathy and atrial fibrillation, where disruption of cell–cell electrical coupling contributes to disease. The DL set also enriches blood microparticle, consistent with its broader capture of platelet- and blood-related biology noted above.

#### KEGG Pathways

Cytoskeleton in muscle cells is the most strongly enriched pathway in both gene sets, together with dilated and hypertrophic cardiomyopathy, directly matching the corresponding disease classes in the benchmark. The ML set additionally enriches fluid shear stress and atherosclerosis, cardiac muscle contraction, and regulation of actin cytoskeleton, pointing toward the aortic and vascular remodeling component of the cohort, whereas the DL set instead enriches glycolysis/gluconeogenesis, the HIF-1 signaling pathway, and ferroptosis, pathways associated with hypoxia and metabolic failure that are well documented in ischemic and infarction-related myocardial injury. The DL set also enriches platelet activation as a KEGG term, reinforcing the thrombotic signal already observed in the biological process results. Overall, the ML-derived biomarkers emphasize structural, contractile, and vascular remodeling mechanisms most relevant to the cardiomyopathy and aortic/ischemic disease groups, while the DL-derived biomarkers additionally highlight platelet-mediated, metabolic, and hypoxia-associated mechanisms relevant to the thrombotic and ischemic disease groups. This divergence indicates that the two learning paradigms capture partially non-overlapping but biologically complementary aspects of cardiovascular pathology.

#### Gene–Pathway Interaction Network

To visualize how individual genes connect to these enriched pathways, a combined KEGG gene–pathway interaction network is constructed from the ML and DL consensus biomarker sets using Cytoscape (Fig. 8). Genes such as ACTG1, TPM3, TTN, DES, ATP2A2, and GNAS are linked to the dilated, hypertrophic, and arrhythmogenic right ventricular cardiomyopathy pathways, consistent with their prioritization as biomarkers for these specific cardiomyopathy classes in Table 2. A larger set of genes, including ACTA1, MYH9, FN1, VIM, ACTB, TLN1, DCN, and DES, connects to the cytoskeleton in muscle cells pathway, underscoring the centrality of cytoskeletal integrity across nearly all cardiac disease classes examined. Genes linked to fluid shear stress and atherosclerosis (FOS, HSP90AB1, DUSP1, ACTG1) point to endothelial and vascular stress mechanisms relevant to the aortic disease group, while genes connected to ferroptosis (FTH1, FTL, PCBP2, ACSL1) implicate iron-dependent oxidative cell death, a mechanism plausibly relevant to ischemic and infarction-related tissue injury. Finally, genes linked to the HIF-1 signaling pathway (ENO1, GAPDH, LDHB, RPS6, ALDOA) suggest hypoxia-driven metabolic adaptation, consistent with the ischemic disease group. Together, the network indicates that the prioritized biomarkers participate in interconnected pathways spanning cardiac structure, vascular stress, oxidative injury, and metabolic regulation, providing a mechanistic complement to the disease-level biomarker associations summarized in Table 2.

**Figure 8:**
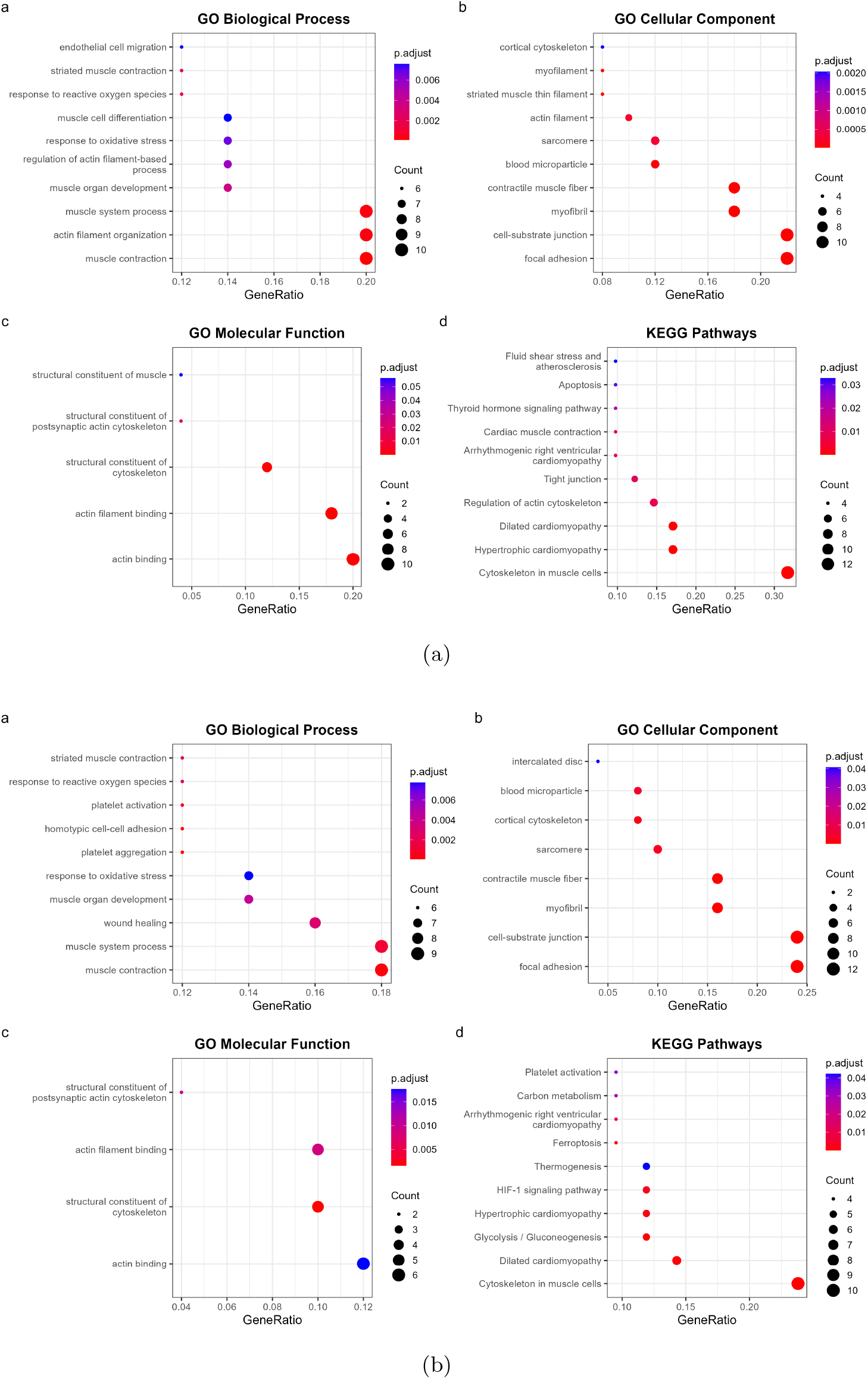
Functional enrichment analysis of consensus biomarkers identified using **(A)** ML and **(B)** DL frameworks. Each subplot includes (a) GO Biological Process, (b) GO Cellular Component, (c) GO Molecular Function, and (d) KEGG pathway enrichment. Dot size represents gene count and color indicates adjusted p-value.

The KEGG pathway enrichment analysis further identified several pathways associated with cardiovascular disease mechanisms. In both ML and DL biomarker sets, significant enrichment is observed in dilated cardiomyopathy, hypertrophic cardiomyopathy, arrhythmogenic right ventricular cardiomyopathy, and cytoskeleton in muscle cells, indicating a potential association between the identified genes and structural cardiac remodeling. The ML-derived biomarkers additionally demonstrated enrichment in fluid shear stress and atherosclerosis, cardiac muscle contraction, regulation of tight junction, actin cytoskeleton, and apoptosis, suggesting stronger involvement in cardiac contractility, vascular remodeling, and endothelial-related mechanisms. In contrast, the DL-derived biomarkers showed enrichment in glycolysis/gluconeogenesis, HIF-1 signaling pathway, carbon metabolism, thermogenesis, ferroptosis, and platelet activation, indicating potential contributions of metabolic dysregulation, oxidative stress, hypoxia-associated pathways, and thrombosis in CVDs development and progression.

Overall, the enrichment analyses demonstrated that both frameworks identified biologically relevant pathways associated with CVDs while also capturing distinct molecular mechanisms. ML-derived biomarkers appeared to emphasize structural and vascular remodeling processes, whereas DL-derived biomarkers additionally highlighted metabolic, hypoxic, and platelet-associated pathways, suggesting complementary biological insights into CVDs pathology.

The KEGG gene–pathway interaction network (Fig. 9), generated using Cytoscape, illustrates the relationship between prioritized genes and significantly enriched pathways potentially associated with CVDs-related mechanisms. The network is constructed by integrating significantly enriched KEGG pathways identified from both the ML and DL consensus biomarker gene sets, thereby providing a combined overview of complementary and shared molecular interactions. Several pathways demonstrated interactions with multiple genes, highlighting interconnected biological processes underlying cardiovascular dysfunction. Notably, pathways related to dilated cardiomyopathy, hypertrophic cardiomyopathy, and arrhythmogenic right ventricular cardiomyopathy are enriched and interconnected with genes involved in cardiac structure, contractility, and myocardial remodeling. Genes such as ACTG1, TPM3, TTN, DES, ATP2A2, and GNAS are linked to these cardiomyopathy-related pathways, suggesting their potential role in maintaining myocardial integrity and cardiac muscle function. A substantial proportion of genes are also associated with the cytoskeleton in muscle cells pathway, including ACTA1, MYH9, FN1, VIM, ACTB, TLN1, DCN, and DES, indicating the importance of cytoskeletal organization and structural stability in cardiovascular pathology. These results support the biological relevance of the identified biomarkers in CVDs progression, since cytoskeletal dysregulation contributes to impaired cardiac contraction and tissue remodeling. In addition, enrichment is observed in pathways associated with atherosclerosis and fluid shear stress, involving genes such as FOS, HSP90AB1, DUSP1, and ACTG1, suggesting a possible contribution to endothelial dysfunction and vascular stress responses. The network further identified genes linked to ferroptosis, including FTH1, FTL, PCBP2, and ACSL1, highlighting mechanisms related to oxidative stress and iron-dependent cell death that may contribute to cardiovascular injury. Moreover, the HIF-1 signaling pathway, involving ENO1, GAPDH, LDHB, RPS6, and ALDOA, suggests potential alterations in hypoxia-associated metabolic adaptation. Collectively, these findings indicate that the prioritized genes participate in diverse but interconnected biological pathways relevant to cardiovascular structure, oxidative stress, metabolism, and disease progression.

**Figure 9:**
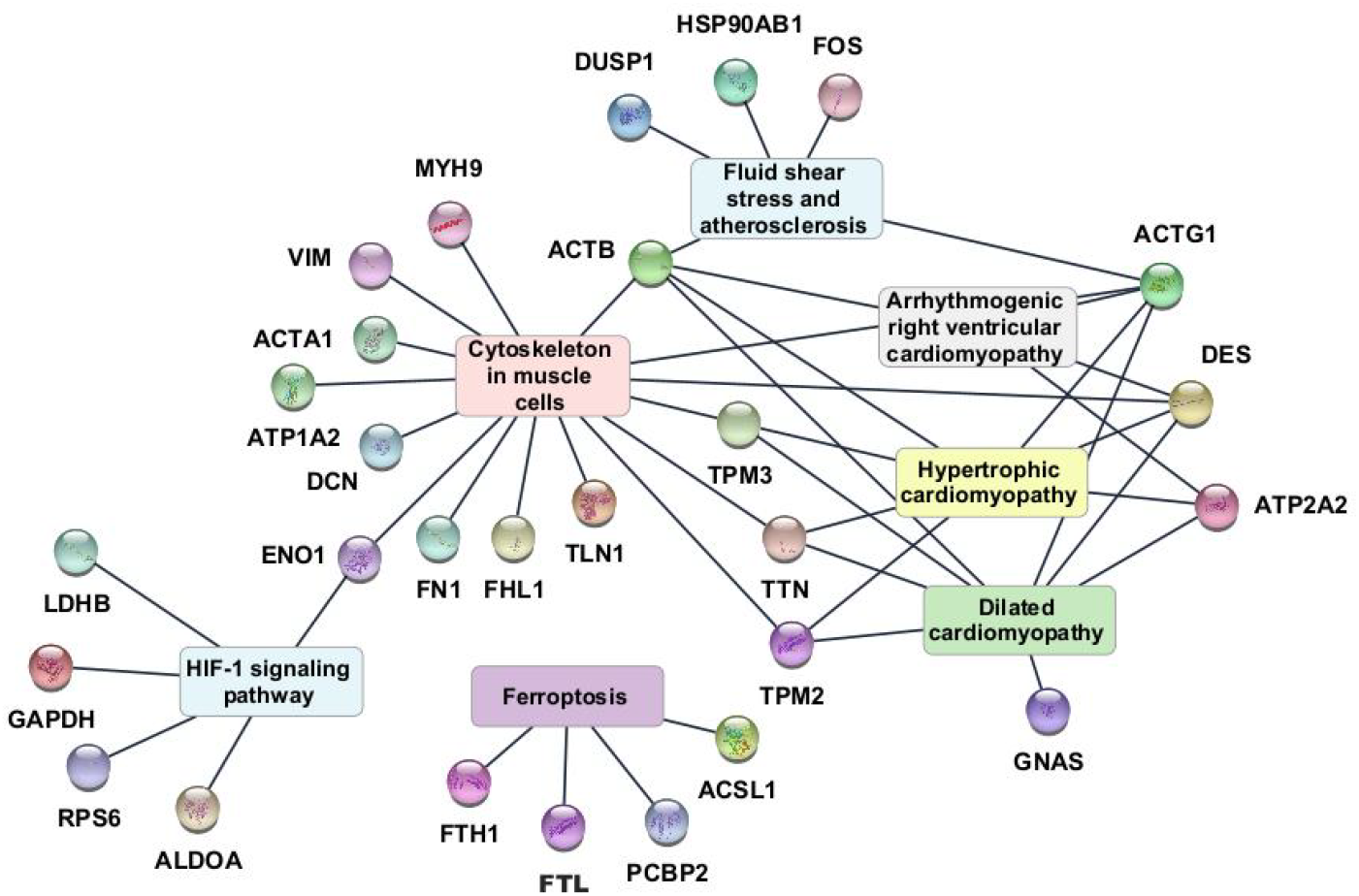
KEGG Pathway Enrichment cnetplot showing the association between enriched pathways and genes. Nodes represent genes and pathways, and edges indicate their interactions.

**Figure 10:**
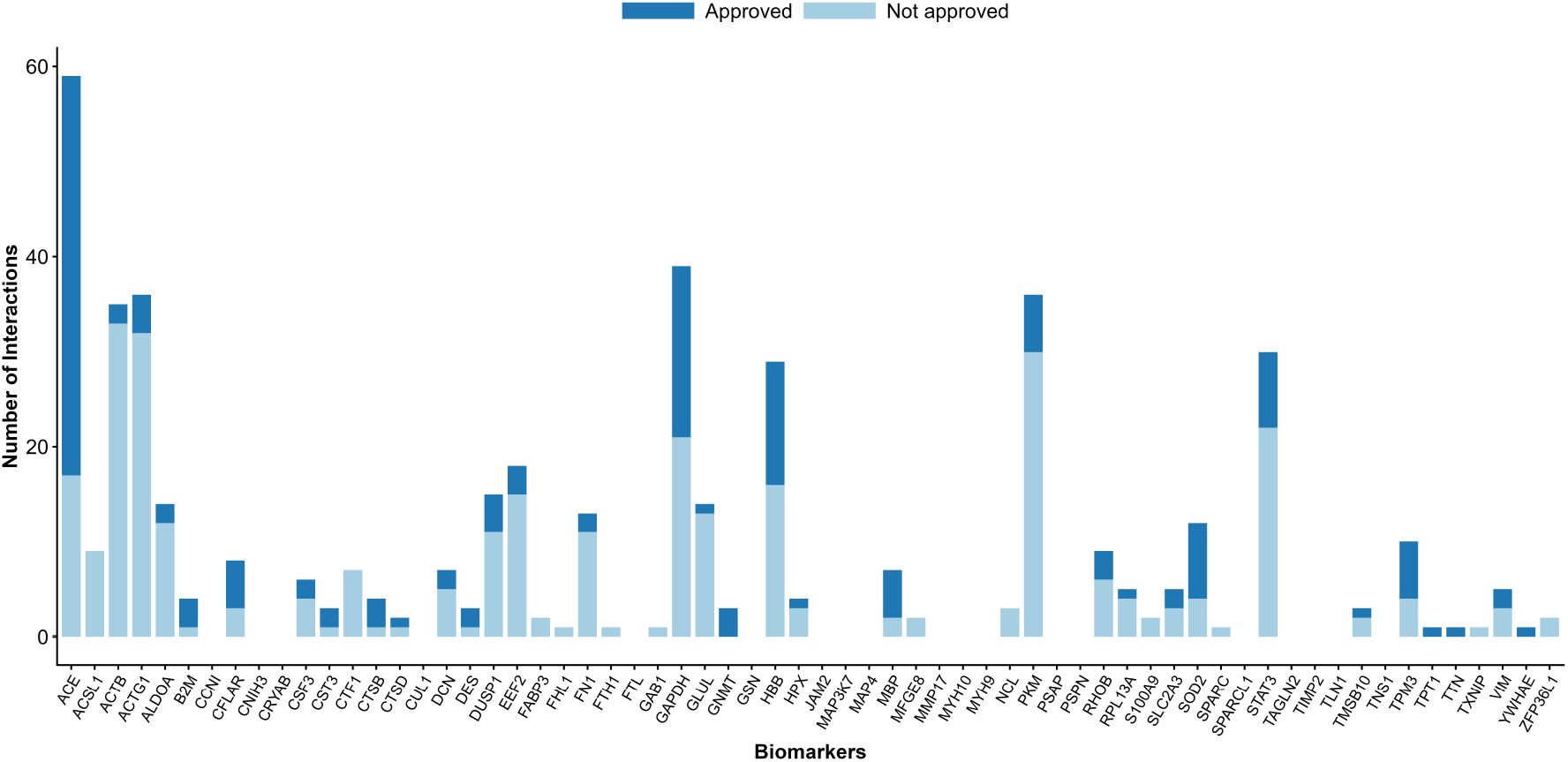
Drug–gene interaction profile of potential CVDs biomarkers. Stacked bars represent the number of approved and non-approved drug interactions reported for each prioritized gene based on DGIdb.

### 2.6 Druggability Analysis

To investigate the therapeutic potential of the identified biomarkers, a druggability analysis is performed using the Drug–Gene Interaction Database (DGIdb), summarized in Table 4. As multiple genes are shared across different cardiovascular disease (CVDs) classes, the potential biomarkers are consolidated into a non-redundant set of unique genes for subsequent analysis. This unique biomarker set is queried against DGIdb to identify known drug–gene interactions.

**Table 4:**
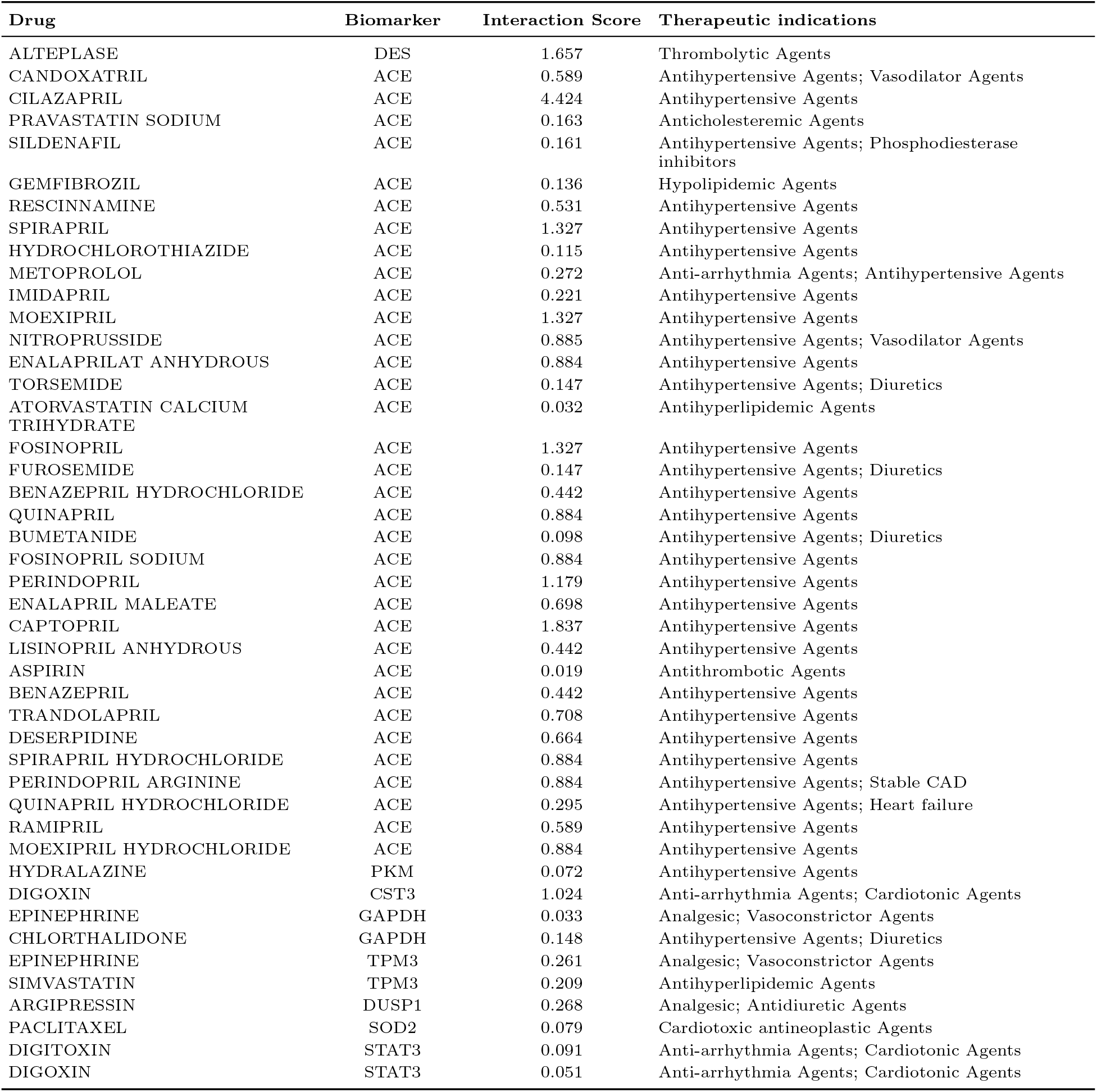
Druggability analysis of prioritized consensus biomarker genes using DGIdb.

The DGIdb search reveals substantial variation in the number of reported drug–gene interactions across the prioritized biomarkers. The highest number of interactions are shown for ACE with 59 (42 approved and 17 non-approved) drug interactions. Other biomarkers with relatively high numbers of interactions are GAPDH (39 interactions, 18 approved), ACTG1 (36 interactions, 4 approved), PKM (36 interactions, 6 approved), ACTB (35 interactions, 2 approved), and STAT3 (30 interactions, 8 approved). Several other biomarkers, such as EEF2, HBB, DUSP1, ALDOA, TPM3, MBP, CFLAR, SOD2, DES and CST3, reported one or more approved drug interactions, which suggests that there are clinically approved therapeutic associations for further evaluation. In contrast, some prioritized biomarkers show limited or no reported drug associations. For the biomarkers: TIMP2, GSN, PSAP, MYH9, MAP3K7, MMP17, TLN1, CNIH3, GSN, PSPN, TNS1, FTL, MAP4, TAGLN2, JAM2, SPARCL1, CUL1, MYH10, or CCNI, no drug interactions are reported in DGIdb. In addition, the following genes are correlated with non-approved drug interactions: MFGE8, SPARC, FHL1, TXNIP, ACSL1, CTF1, NCL, FABP3, GAB1, S100A9, ZFP36L1, and FTH1, suggesting that no approved therapeutic associations are currently available for these biomarkers in DGIdb. To further assess the clinical relevance of the analysis, the approved drug–gene interactions identified from DGIdb are evaluated based on their reported therapeutic indications. Since approved drugs may not necessarily be indicated for CVDs, the therapeutic indications of all approved drugs are subsequently verified using DrugBank, and only drugs with established cardiovascular-related indications are retained. Based on this filtering strategy, DES, ACE, PKM, CST3, GAPDH, TPM3, DUSP1, SOD2, and STAT3 are identified as biomarkers with approved cardiovascular-relevant drug interactions and are therefore selected for detailed analysis in the subsequent section. The curated cardiovascular drug–gene interactions, together with their DGIdb interaction scores and therapeutic indications, are summarized in Table 3. Among the retained interactions, ACE exhibits the widest range of approved therapeutics for CVDs, highlighting its established place as a therapeutic target for cardiovascular disease management. The majority of ACE-associated drugs belong to the antihypertensive class, including cilazapril, spirapril, imidapril, moexipril, fosinopril, quinapril, perindopril, enalapril, captopril, lisinopril, ramipril, trandolapril, benazepril, and related formulations. Additional cardiovascular therapies targeting ACE included vasodilators (candoxatril and nitroprusside), diuretics (hydrochlorothiazide, torsemide, and bumetanide), lipid-lowering agents (pravastatin, atorvastatin, and gemfibrozil), the anti-arrhythmic agent metoprolol, the antithrombotic agent aspirin, and sildenafil, a phosphodiesterase inhibitor with approved cardiovascular applications. Collectively, these interactions demonstrate the extensive therapeutic landscape associated with ACE across multiple cardiovascular treatment strategies. In addition, a number of other biomarkers show approved drug interaction for different cardiovascular therapeutic classes. Alteplase, a thrombolytic agent used in the management of acute thrombotic events is associated with DES. The associations of PKM with an antihypertensive agent, hydralazine, whereas CST3 and STAT3 are associated with the cardiotonic and anti-arrhythmic agents digoxin and digitoxin, respectively. GAPDH and TPM3 show interactions with drugs representing different therapeutic classes in CVDs including the lipid-lowering agent simvastatin, the vasoactive agent epinephrine, and the diuretic chlorthalidone, which is used in cardiovascular emergency settings. DUSP1 is found to be associated with the antidiuretic and vasoactive agent, argipressin, which is used in cardiovascular applications in critical care, while SOD2 is associated with paclitaxel, a drug with well established cardiovascular relevance, though classified as an antineoplastic agent, used in the development of vascular drug-eluting devices.

## 3 Discussion

In this research, an XAI framework for CVDs is presented, and a workflow is proposed that not only considers predictive performance but also evaluates various XAI methods systematically to ensure that the attribution of biomarkers is reliable, prioritizing those related to the disease. The sequential approach enables robust identification of genes of interest for biological relevant pathways without losing interpretability along the analytical process.

The quality of omics data integration is a key determinant in downstream analyses, especially if the data is from different studies, tissues, and experimental platforms. Across all evaluated datasets, Shambhala-2 consistently demonstrated superior performance for RNA and combined RNA/MH datasets, whereas all three harmonization methods produced comparable improvements for MH datasets. These results indicate that harmonization can be more effective depending on the molecular modality of the underlying data; the Shambhala-2 transformation seems to be more effective on transcriptomic data. Selecting the harmonization strategy based on quantitative evaluation, rather than applying a single preprocessing approach indiscriminately, strengthens the reliability of subsequent analyses by minimizing the influence of platform-specific variability on disease classification and biomarker discovery.

The improvements achieved during harmonization are reflected in the classification performance obtained across multiple ML and DL algorithms. The strong performance observed across both paradigms indicates that the harmonized data possessed robust disease associated molecular patterns and effectively reduced technical variability. Ensemble learning algorithms are well suited to the high dimensional nature of omics data, as they are able to model complex nonlinear relationships and are often less sensitive to residual data heterogeneity, while neural network-based architectures offer the ability to learn hierarchical feature representations to capture subtle biological properties across a variety of molecular profiles. The ability of these fundamentally different learning paradigms to achieve consistently high predictive performance indicates that the identified disease signatures are not dependent on a specific modeling strategy, thereby providing a reliable foundation for subsequent explainability analyses and consensus-based biomarker prioritization.

Although prediction is crucial, the acceptance of AI in the clinical field relies heavily on the knowledge of how such models make decisions. Feature importance scores vary depending on the method of explainability, which results from a different mechanism of calculating the importance of features and consequently, it can yield significantly different explanations for the same prediction. This highlights the importance of objectively evaluating explanation quality before using XAI-derived features for biomarker discovery. The proposed framework reduces model-specific bias by choosing the best valid explanation method applied to each classifier and combines the selected genes by consensus ranking, thus ensuring that prioritized biomarkers represent reproducible molecular signatures rather than artifacts of individual prediction or attribution algorithms.

The functional enrichment analysis showed that the prioritized consensus biomarkers are involved in biological processes and signaling pathways that are well characterized in the pathogenesis of cardiovascular disease, which suggested the biological validity of the proposed biomarker discovery framework. The ML and DL derived gene sets showed different patterns of enrichment, but both shared consistent enrichments of complementary mechanisms involving cardiac function, structural remodeling, metabolic regulation, and oxidative stress. The convergence of these learning paradigms indicates that by combining biomarkers from various predictive models, one can gain insights into the various interconnected aspects of cardiovascular pathology. Together, these findings suggest that a multimodal integration of complementary computing methods will benefit biological interpretation of complex diseases such as CVDs, and lead to a more comprehensive molecular understanding of cardiovascular disease mechanisms.

Consensus-based biomarker prioritization combined with functional characterization and druggability provides an additional dimension of translational relevance beyond accurate disease classification for the proposed framework. The presence of both common and disease-specific biomarkers indicates that CVDs are a complex disease group, with common mechanisms despite the specific nature of the disease. The strategy supports the use of biomarkers consistently supported by a variety of high-performing predictive models and explainability methods, thereby enhancing the confidence in the biological relevance of the identified molecular signatures and minimizing reliance on specific computational models. In addition, the fact that several prioritized biomarkers are linked to already approved cardiovascular drugs highlights the clinical relevance of the framework and its potential to support drug repurposing and biomarker validation and therapeutic target prioritization. Furthermore, it is likely that biomarkers with no known cardiovascular drug associations could serve as potential targets for future functional studies, highlighting the importance of combining XAI with subsequent downstream biological and pharmacological studies, bringing the computational discovery of biomarkers closer to the realm of translational cardiovascular studies.

Despite these encouraging findings, several limitations should be acknowledged. First, the analyses are done with publicly available datasets from various studies, in which biological and technical heterogeneity can still arise from differences in patient characteristics, sample acquisition protocols, and experimental platform, despite the harmonization. Second, biomarker prioritization is done exclusively by computational analyses using XAI and therefore requires independent experimental validation before biological or clinical conclusions can be established. Third, while the consensus ranking approach mitigates the reliance on any particular model and explanation method, the top-ranked biomarkers are still subject to the classifiers selected and explainability methods assessed in this work. Further studies with other predictive architectures and newly developed approaches to XAI may further improve biomarker robustness. Last, the druggability analysis relied on the available databases of drug-gene interactions and validated therapeutic indications, which provide evidence of drug-gene interaction but are not proof of therapeutic efficacy in the CVDs studied here.

Future studies should thus examine the prioritized biomarkers in independent patient cohorts with wider demographic and clinical diversity to confirm their results. Experimental studies that explore the functional roles of these identified genes will be crucial in elucidating their mechanistic involvement in CVDs pathogenesis. The incorporation of other omics layers such as proteomics, metabolomics, and single-cell sequencing could further enhance molecular characterization and uncover complementary markers not detectable using transcriptomics. Further, prospective clinical trials testing the diagnostic, prognostic, and therapeutic value of the biomarkers identified will be necessary before these findings can be translated into routine clinical practice. Refined XAI approaches and extensive biological validation will further support development of accurate and clinically actionable computational tools for identifying cardiovascular biomarkers.

## Notes

### Competing Interest Statement

The authors have declared no competing interest.

